# Combined single-cell and spatial transcriptomics reveals the molecular, cellular and spatial bone marrow niche organization

**DOI:** 10.1101/718395

**Authors:** Chiara Baccin, Jude Al-Sabah, Lars Velten, Patrick M. Helbling, Florian Grünschläger, Pablo Hernández-Malmierca, César Nombela-Arrieta, Lars M. Steinmetz, Andreas Trumpp, Simon Haas

## Abstract

The bone marrow (BM) constitutes the primary site for life-long blood production and skeletal regeneration. However, its cellular composition and the spatial organization into distinct ‘niches’ remains controversial. Here, we combine single-cell and spatially resolved transcriptomics to systematically map the molecular and cellular composition of the endosteal, sinusoidal, and arteriolar BM niches. This allowed us to transcriptionally profile all major BM resident cell types, determine their localization, and clarify the cellular and spatial sources of key growth factors and cytokines. Our data demonstrate that previously unrecognized Cxcl12-abundant reticular (CAR) cell subsets (i.e. Adipo- and Osteo-CAR cells) differentially localize to sinusoidal or arteriolar surfaces, locally act as ‘professional cytokine secreting cells’, and thereby establish distinct peri-vascular micro-niches. Importantly, we also demonstrate that the 3-dimensional organization of the BM can be accurately inferred from single-cell gene expression data using the newly developed RNA-Magnet algorithm. Together, our study reveals the cellular and spatial organization of BM niches, and offers a novel strategy to dissect the complex organization of whole organs in a systematic manner.

**One Sentence Summary:** Integration of single-cell and spatial transcriptomics reveals the molecular, cellular and spatial organization of bone marrow niches

## INTRODUCTION

Bone marrow (BM) niches are specialized microenvironments where distinct mesenchymal cells, the vasculature, nerve fibers and differentiated hematopoietic cells interact to govern the maintenance and differentiation of hematopoietic and mesenchymal stem cells^1–3^. Classically, BM niches have been studied by genetic approaches that involve labeling cell types and deleting candidate niche factors based on the expression of individual reporter genes. These studies have generated important insights into the functional roles and cellular sources of key cytokines such as Cxcl12 or Scf (*Kitl*)^4–10^. However, in such approaches a single marker is used to define cell types, likely resulting in the labelling of heterogeneous populations instead of specific cell types. These short-comings have resulted in a controversial debate about the importance of distinct cell types and factors, and their localization to peri-sinusoidal, peri-arteriolar or endosteal niches^4–10^. To gain a global understanding of cell types and niches in the BM, we have generated a single-cell RNA-sequencing (scRNAseq)-based molecular map of all major BM populations. We then used spatially resolved transcriptomics in combination with novel computational tools to allocate cell types to different BM niches, determine molecular mediators of intercellular interactions, and identify the cellular and spatial sources of niche factors.

## RESULTS

### Identification and characterization of BM resident cell types by scRNAseq

Frequencies of BM cell types differ by several orders of magnitude^11^, imposing a challenge to scRNAseq approaches. To capture both highly abundant as well as extremely rare BM resident cells, we performed droplet-based scRNAseq^12^ of cells from total mouse BM, followed by progressive depletion of highly abundant cell types or enrichment of rare populations from undigested BM or enzymatically digested bones (Figure 1a). In total, our dataset comprises 7497 cells with a median detection of 1999 genes per single cell, which formed 32 clusters corresponding to distinct cell types or stages of differentiation (Figure 1b, S1). Importantly, this map is not quantitative with regard to the relative size of the different cell populations, since dissociation rates largely differ between cell types^11^. As detailed below, cell type annotation was performed based on marker gene expression (Table S1, Figure S2,3), gene ontology analyses (Table S1), and by quantifying the enrichment of cluster gene signatures in transcriptomic data of previously described bulk populations using the CIBERSORT algorithm^13,14^ (Figure S4, Supplementary Note). We used SOUP^15^ to confirm that the mesenchymal populations described in the following are primarily distinct cell types, while transition states between clusters exist (Figure S5).

**Figure 1.**
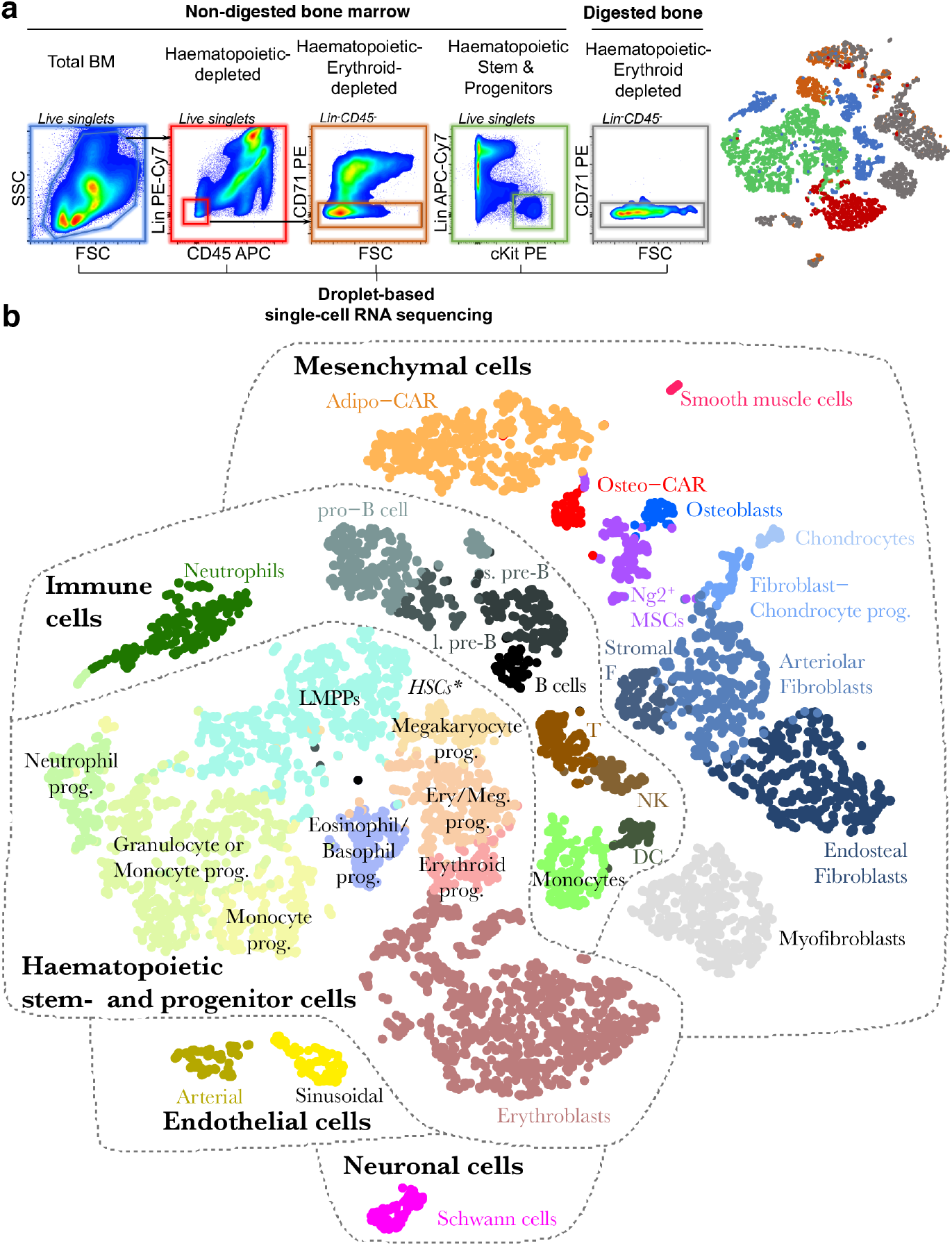
Identification of BM resident cell types by scRNA-seq. **a**, Overview of the FACS sorting strategy. In total, 5 consecutive single-cell RNA sequencing runs were performed. Right panel: t-SNE projection of all cells with respective experiment colour-coded. **b**, t-SNE projection of all cells with clusters colour-coded. Abbreviations used: T: T cells, NK: Natural killer cells, s. pre-B: small pre-B cells, l. pre-B: large pre-B cells, DC: Dendritic cells, Prog.: progenitor. * Lin^−^Kit^+^Sca1^high^ HSCs reside at the interface of LMPPs and Megakaryocyte progenitors, see also figure S3g.

As expected, scRNAseq of total bone marrow identified the major haematopoietic and immune cell types, including dendritic cells, neutrophils, monocytes, T cells, and distinct developmental B cell stages (Figure 1b, S2, S4a). Upon depletion of these major immune populations, scRNAseq primarily yielded erythroid progenitors that exhibited low expression of the pan-haematopoietic marker CD45 and displayed erythroid markers such as CD71^16^ (Figure S2). An additional 2% of these cells were non-haematopoietic (Figure 1a, S1). To efficiently capture non-haematopoietic cells in depth, we subsequently depleted cells expressing lineage markers, CD45 or CD71 from non-digested bone marrow and enzymatically digested bone chips (Figure 1a). This allowed us to interrogate rare cell populations, including neural Schwann cells (*Mog*, *Mag*), smooth muscle cells (*Tagln*, *Acta2*), putative myofibroblasts, nine different *Pdgfra*-positive mesenchymal cell populations and two endothelial cell clusters (*Cdh5*, *Pecam*) (Figure S3, Table S1 for a list of signature genes). The endothelial populations comprised Sca-1 (*Ly6a*)-expressing arterial and *Emcn*-expressing sinusoidal endothelial cells, respectively (Figure 2a, b)^17^. Among the mesenchymal cell populations, we identified chondrocytes (*Acan*, *Sox9*), osteoblasts (*Osteocalcin/Bglap*, *Col1a1*), as well as several less well described cell types. First, we observe three distinct fibroblast-like populations, which we further annotate based on their differential localization below. Second, we observe two additional populations that showed high transcriptomic similarity to previously described SCF-GFP^+5^ and Cxcl12-GFP^+9^ cells, also called Cxcl12-abundant reticular (CAR) cells (Figure 2c)^4,18^. Remarkably, these populations differentially expressed adipocyte and osteo-lineage genes (Figure 2d), while exhibiting similar overall transcriptomic profiles. As a consequence, we termed these previously undescribed subpopulations Adipo- and Osteo-CAR cells, respectively. Adipo-CAR cells expressed high levels of leptin receptor (*Lepr*), and showed high transcriptomic similarity to previously described LepR-Cre cells^5,8,19^ (Figure 2d). In contrast, Osteo-CAR cells expressed higher osterix (Sp7, Figure 2d, S3n) and lower Lepr levels. Third, we identified a cluster of Ng2 and Nestin-expressing mesenchymal cells that show transcriptomic similarity to previously described Ng2^+^Nestin^+^ mesenchymal stem cells (MSCs), which we therefore termed Ng2^+^ MSCs (Figure 2a,c). Ng2^+^Nes^+^ MSCs are clearly distinct from Nes^+^ endothelia^20^ or smooth muscle cells^21^. Pseudotime analysis using RNA-Velocity^22^ placed Ng2^+^ MSCs at the apex of a differentiation hierarchy with osteoblasts, CAR cells, chondrocytes, and fibroblasts being downstream (Figure 2e).

**Figure 2.**
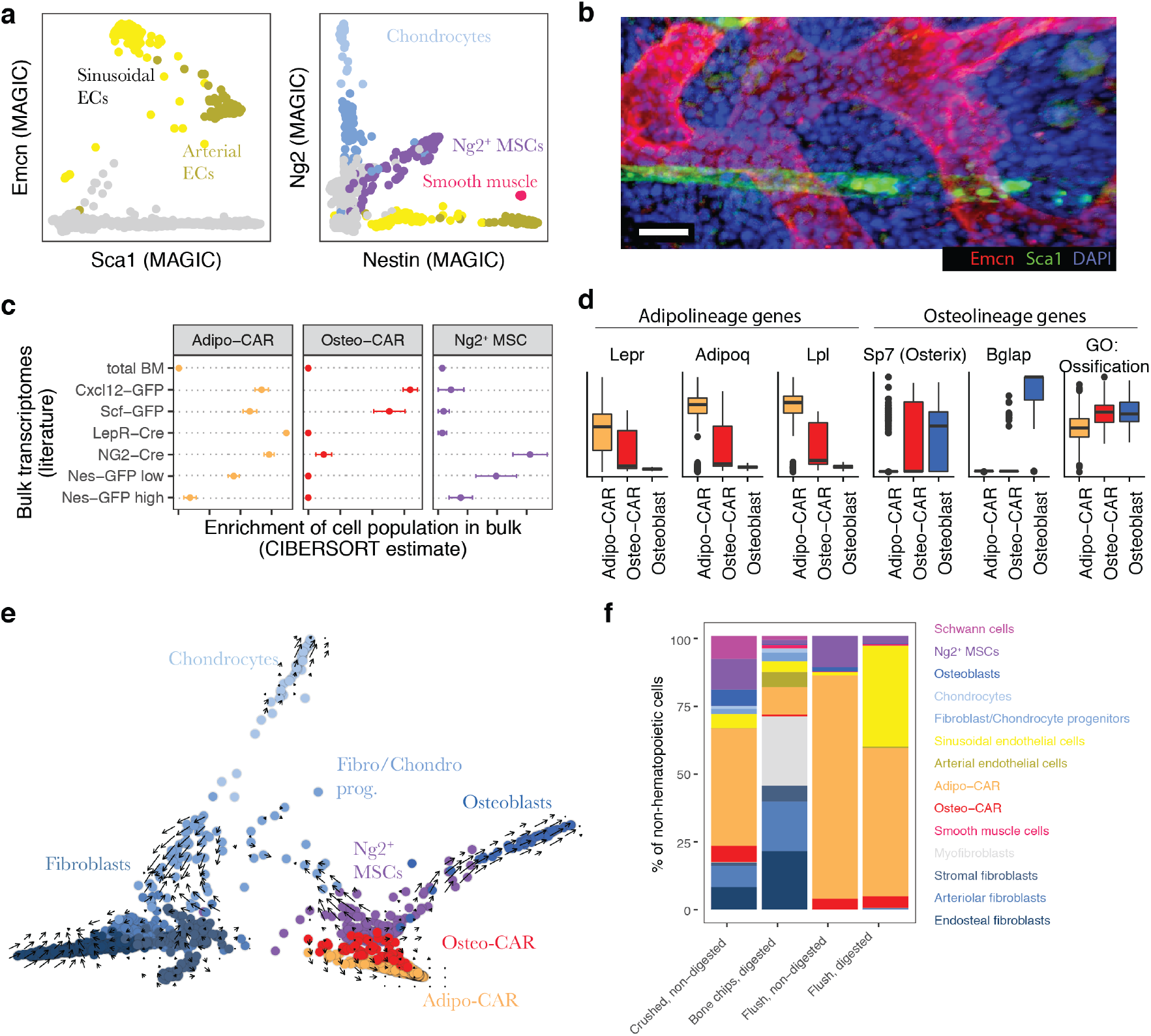
Characterization of BM resident cell types. **a**, Gene expression levels of Sca1, Endomucin (Emcn), Nestin, and Ng2, with relevant populations colour-coded. MAGIC^47^ was used for imputation of drop-out values. **b**, Deep imaging of a BM section immunostained with antibodies against Sca-1 and Emcn. Scale bar = 20 μm. **c**, Enrichment of gene expression signatures of Ng2^+^ MSCs, Adipo-CAR and Osteo-CAR cells in previously published transcriptomes of relevant genetically labelled populations^5,7–9^. See Figure S4b for further populations, and the supplementary note for a detailed evaluation of the algorithm used (CIBERSORT). Error bars indicate the standard error of the mean across n=3 bulk transcriptome samples per class. **d**, Boxplots of the scaled expression level of selected genes in single cells from the Adipo-CAR, Osteo-CAR and Osteoblast populations. For the right panel, the mean expression of all genes annotated with the gene ontology term ‘ossification’ was computed for each cell. **e**, Projection of all mesenchymal cell types using PHATE^48^ with time derivatives of gene expression state, as determined by RNA velocity^22^, highlighted as arrows. **f**, Comparison of cell type frequencies between distinct cell isolation methods used for scRNAseq.

To comprehensively cover haematopoietic stem and progenitor cell (HSPC) populations, we additionally performed scRNAseq of Lin^−^cKit^+^ cells. This revealed a differentiation continuum^23–26^ spanning megakaryocyte-erythrocyte and lympho-myeloid branches, as well as a separate cluster of eosinophil/basophils progenitors (Figure 1b, S2).

In order to investigate the impact of the isolation strategies of BM resident cells on the cell type recovery in scRNAseq experiments, we compared flushing of BM versus crushing of whole bones to release the marrow, and evaluated the impact of enzymatic digestion. This demonstrated that the majority of cell populations are found both in flushed and crushed BM, but many populations are only released efficiently upon intense physical treatment or enzymatical digestion (Figure S6, Figure 2f). In particular, the fibroblast populations were found more abundantly in crushed if compared to flushed bones. While we have confirmed the presence of these subpopulations in the diaphyseal BM using imaging and spatial transcriptomics (see Figure 3, 4c-f), similar cell types deriving from the cortical bone, epiphysis or periosteum might also be present in scRNAseq datasets from whole bones. Only the myofibroblast and Schwann cell populations were exclusively present in the crushed bone samples, suggesting that they are either firmly attached to the bones or may derive from distal regions, such as the periosteum or the epiphysis.

**Figure 3.**
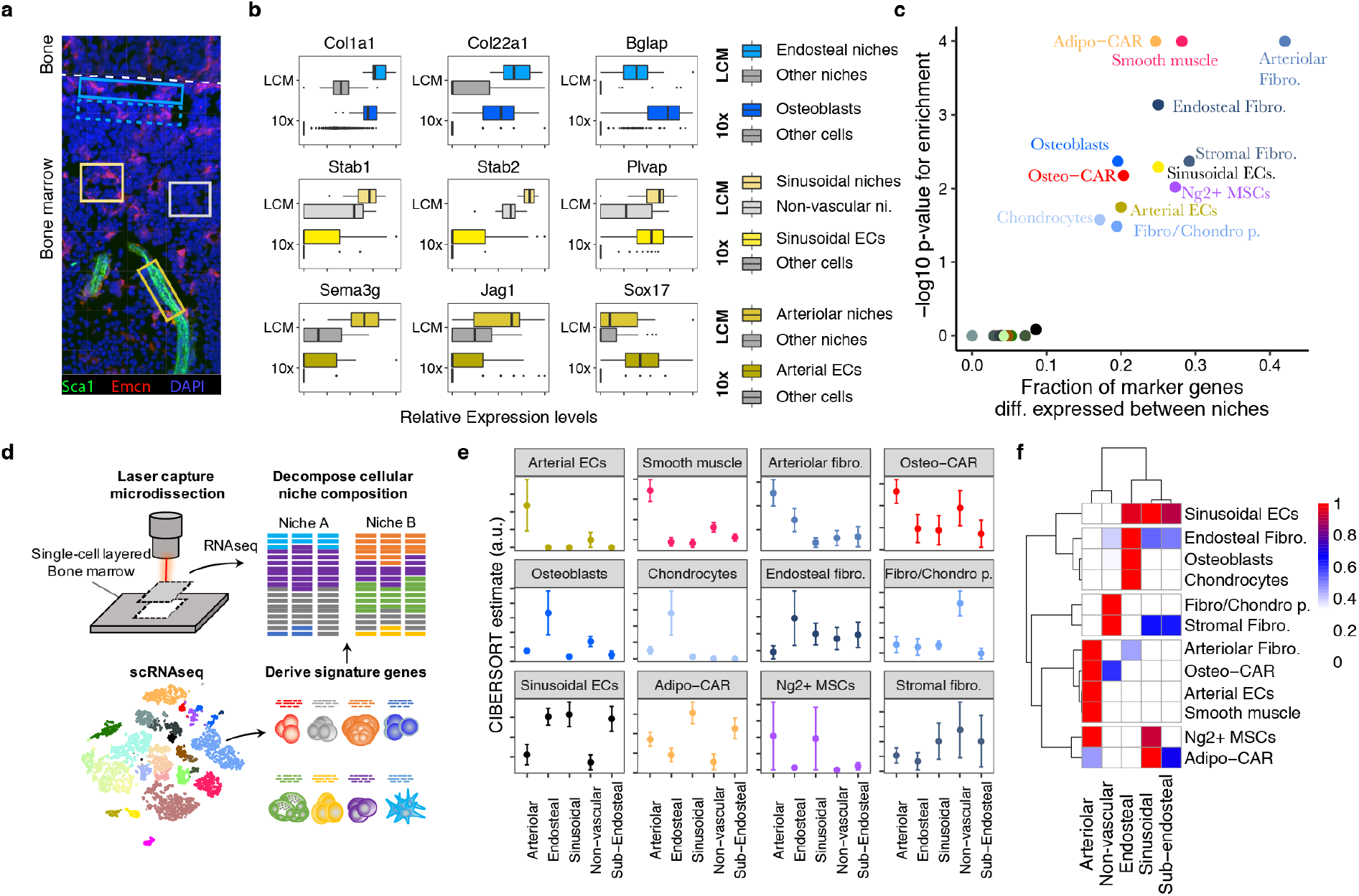
Spatial allocation of BM resident cell types by integrated single-cell and spatial transcriptomics. **a**, Scheme of experimental design. 12 μm bone sections were stained for arterioles (Sca1) and sinusoids (Emcn), respectively. Areas of approximately 14.500 μm^2^ surrounding arteries (dark yellow box), sinusoids (yellow box) and the endosteum (blue box), as well as areas with no vessels (grey box) and sub-endosteal areas (dotted blue box) were collected by laser capture dissection and subjected to RNA-seq. A confocal image is shown for illustrative purposes. For images acquired under the laser capture dissection microscope and selected areas see Figure S7b. **b**, Expression of osteoblast-, sinusoid- and arteriole-specific genes in scRNA-seq data (10x) and spatial transcriptomics from different niches (LCM: LCM-seq data). **c**, Enrichment of population marker genes (Table S1) among genes with differential expression between niches (Table S2). **d**, Schematic outline of the computational data analysis strategy used. **e**,**f**, Estimated abundance of different cell types in microscopically distinct niches. Error bars indicate standard error of the mean of the estimate across n=11 to n=28 samples per class.

**Figure 4.**
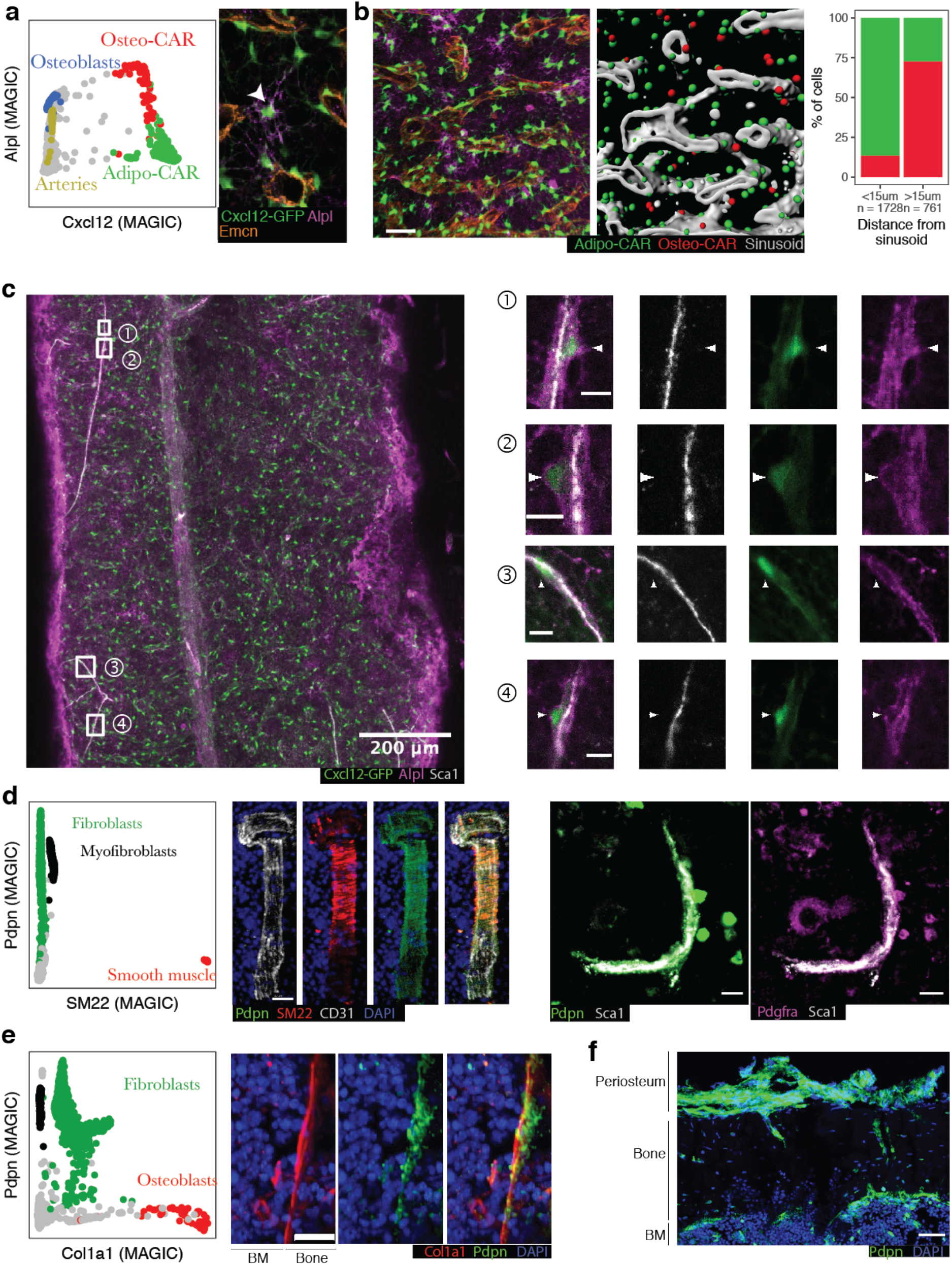
Localization of key mesenchymal cell types by immunofluorescence. **a**, Left panel: Single-cell gene expression levels of *Cxcl12* and *Alpl* with relevant cluster identity colour-coded. MAGIC^47^ was used to impute drop-out values. Right panel: Sample high-resolution ROI from a whole-mount image of a femur from a Cxcl12-GFP mouse, stained for Alpl and the sinusoidal marker Emcn. An Alpl^+^Cxcl12-GFP^+^ cell distant from sinusoids is highlighted by an arrowhead. **b**, Quantitative analysis of a full whole-mount image, see also Figure S9a. Left panel: Sample ROI, scale bar: 50μm. Central panel: 3D segmentation of the same ROI. Cxcl12-GFP^+^ cells were classified as Osteo- or Adipo-CAR cells based on the Alpl signal. Right panel: Quantitative assessment of Alpl^+^ Osteo-CAR cells (red) and Alpl^neg^ Adipo-CAR cells (green) between sinusoidal and non-sinusoidal niches in central BM. **c,** Whole-mount imaging of a femur from a Cxcl12-GFP mouse, stained for Alpl and the arteriolar marker Sca1. Arrowheads point to Alpl^+^Cxcl12^+^ cells near, but not overlapping with, Sca1^+^ arteriolar endothelial cells. Scale bars in ROIs: 10μm. See figure S9b for a second whole-mount image. **d,** Left panel: Single-cell gene expression levels of SM22 (*Tagln*) and *Pdpn* with relevant cluster identity colour-coded. Central panel: Immunofluorescence staining of a BM arteriole stained for SM22, Pdpn and CD31. Right panel: Immunofluorescence staining of a BM arteriole stained for Pdpn, Sca1 and Pdgfra. Scale bar: 20 μm. **e,** Left panel: Single-cell gene expression levels of *Col1a1* and *Pdpn* with relevant cluster identity colour-coded. Right panel: Immunofluorescence staining of Col1a1, Pdpn and DAPI at the endosteal surface. Scale bar: 20 μm. **f**, Immunofluorescence staining of Pdpn at the endoestum, cortical bone and periosteum.

In summary, our dataset constitutes the most comprehensive scRNAseq study of the homeostatic BM to date^21,27^ (Figure S6e), and spans almost all BM resident cell types described previously as well as several novel cell types. Osteoclasts, neurons and mature megakaryocytes are not covered by our dataset, likely due to cell size limitation of the scRNAseq approach. The full dataset can be interactively browsed at http://nicheview.shiny.embl.de

### Spatial allocation of BM resident cell types by combined single-cell and spatial transcriptomics

While single-cell transcriptional profiling provides a powerful tool to characterize the identity and molecular makeup of BM resident cell types, information about their spatial distribution is lost in such experiments. To gain spatial information on the cell types identified above, we considered integrating our scRNAseq dataset with spatially resolved transcriptomics data of the BM. Recently described spatial transcriptomic approaches require high image and RNA quality^28–30^ or rely on unfixed tissue material^31,32^, and have therefore not been successfully adapted to the adult BM. Here we developed a robust and significantly improved version of laser-capture microdissection-sequencing (LCM-seq)^33^, based on random priming that copes with fixed material and low input RNA quality, and is therefore compatible with fixed bone marrow sections (Figure S7a, see methods). Using this approach, we generated full-length, high quality transcriptomic data from LCM-dissected areas of fixed BM sections containing 200-300 cells in a single cell layer. We applied LCM-seq to 78 microdissected regions collected from the central diaphyseal bone marrow, based on presence or absence of sinusoidal and arteriolar blood vessels, or based on the distance from the endosteum, in order to characterize the endosteal, sub-endosteal, arteriolar, sinusoidal, and non-vascular niche composition (Figure 3a, S7b). Spatially resolved regions from the endosteum and sub-endosteum were collected solely based on the distance to the bone lining, and independent on the presence of blood vessels. Due to the high vascularization of bone marrow, ‘non-vascular’ niches can be expected to be in close proximity but not directly adjacent to sinusoids^11^.

To evaluate our approach, we selected marker genes of cell types known to be specifically present in the respective niches, and compared their expression in scRNAseq and corresponding spatial transcriptomics data (Figure 3b, Table S2). As expected, osteoblast genes were selectively enriched at the endosteum, genes specific for sinusoidal endothelial cells were enriched at regions with high abundance of sinusoids, and arterial endothelial genes were enriched on arterioles. Marker gene sets for haematopoietic populations, Schwann cells and myofibroblasts were not significantly associated with any of the defined niches, suggesting that these cell types are either relatively evenly distributed across niches (haematopoietic populations), or insufficiently covered in the LCM-seq data (Schwann cells and myofibroblasts) (Figure 3c, Figure S7d,e). In contrast, marker gene expression of the remaining 12 BM resident populations differed significantly across analysed niches.

To systematically assess the preferential localization of these BM resident cell types to candidate niches, we computationally estimated the frequencies of cell populations defined by scRNAseq in the spatially resolved transcriptomics data using the CIBERSORT algorithm ^13,14^ (Figure 3d). We extensively validated the ability of CIBERSORT to decompose bulk transcriptomes using a single-cell reference, and evaluated its performance on assembled pools of 100 cells with known composition that were processed using the LCM-seq protocol (see Supplementary Note and Figure S8). As expected, osteoblasts and chondrocytes were found exclusively at *endosteal niches* (Figure 3e,f). In addition, a specific fibroblast population localized preferentially to the endosteum and was therefore tentatively termed endosteal fibroblasts. In contrast, arterial endothelial cells, smooth muscle cells, and a distinct fibroblast population localized specifically to *arteriolar niches* (Figure 3e,f). According to its localization we tentatively termed this fibroblast population arteriolar fibroblasts. Sinusoidal endothelial cells were found in *sinusoidal niches*, but were also present in regions not selected based on presence of vascular subtypes (i.e. (sub)-endosteal niche), in accordance with a widespread web of sinusoids spanning the entire bone marrow (Figure 2b, S9a). Adipo-CAR cells were also found predominantly in areas with high sinusoidal occurrence, in line with their similarity to LepR-Cre cells and the reported peri-sinusoidal localization of these cell types ^4,5^. In contrast, previously not described Osteo-CAR cells preferentially localized to arteriolar or non-vascular niches, suggesting that the two newly described CAR cell populations occupy distinct vascular niches. Ng2^+^ MSCs could not be unanimously placed to a niche, potentially due to additional heterogeneity of this population (Supplementary Note, Figure S8i). In summary, systematically integrating spatial transcriptomics with single-cell transcriptomics data allowed us to localize the majority of known and newly defined BM resident populations to distinct endosteal, sinusoidal, arteriolar and non-vascular niches.

### Validation of cell type localization

To confirm the spatial relationships of BM cell types identified by LCM-Seq, we determined marker combinations specific to the individual populations and performed immunofluorescence imaging of bone sections.

Gene expression analyses suggested that differential expression of alkaline phosphatase (Alpl) and Cxcl12 permits the discrimination of Osteo-CAR cells (Cxcl12^+^Alpl^+^) and Adipo-CAR cells (Cxcl12^+^Alpl^neg^) (Figure 4a). In contrast, osteoblasts, MSCs and arterial ECs expressed Alpl, but only low levels of Cxcl12 (Cxcl12^neg^Alpl^+^). To confirm the *in situ* localization of Adipo- and Osteo-CAR cells, we performed whole-mount immunofluorescence imaging of long bones from Cxcl12-GFP mice^4^. Co-staining with the sinusoidal maker Emcn revealed that Cxcl12^+^Alpl^neg^ Adipo-CAR cells in the central BM predominantly ensheathed sinuosoids, in line with results from LCM-seq (Figure 4a,b, S9a). In contrast, Cxcl12^+^Alpl^+^ Osteo-CAR cells in central BM typically showed a non-sinusoidal localization and a highly reticular morphology (Figure 4a,b S9a). By co-staining with the arterial maker Sca1, we also found numerous instances of Cxcl12^+^Alpl^+^ Osteo-CAR cells in immediate vicinity of Sca1^+^ arterioles (Figure 4c, S9b). In some instances, these formed highly reticulate structures (Figure S9b). We further noticed that GFP^+^ protrusions from Osteo-CAR cells covered Sca1^+^ arterioles, whereas endothelial cells contributed only faint levels of Cxcl12-GFP (Figure 4c, S9b). Together, these observations confirm the predominantly arteriolar and non-vascular localization of Osteo-CAR cells, as predicted from LCM-seq. Importantly, ubiquitous Alpl staining at the endosteum and sub-endosteum (possibly derived from osteoblasts) prevented analysis of CAR cell populations in these regions.

Next, we used CD31, SM22, Pdpn, Pdgfr, Col1a1 and Sca1 as markers to specifically identify smooth muscle cells (SM22^+^Pdpn^neg^), fibroblast populations (Pdpn^+^Pdfgr^+^), osteoblasts (Pdpn^neg^Col1a1^+^), and arterial endothelial cells (Pdpn^neg^CD31^+^Sca1^+^) to localize them *in situ* (Figure 4d-f). As suggested by LCM-seq, CD31/Sca1-expressing arterioles were enveloped by SM22^+^Pdpn^neg^ smooth muscle cells and Pdpn^+^ arteriolar fibroblasts (Figure 4d). Arteriolar fibroblasts appear to be the cellular source for the collagen layer of the tunica externa surrounding arterioles (Figure S7c) and are likely overlapping with previously described peri-arteriolar Pdpn-expressing stromal cells^34^. Moreover, immunofluorescence confirmed the existence of Pdpn^+^Col1a^low^ fibroblasts localizing to the bone-facing side of the endosteal lining made up of Pdpn^neg^Col1a^high^ osteoblasts (Figure 4e). Notably, Pdpn^+^ cells were also found at the cortical bone and periosteal regions, making it possible that fibroblast-like cells similar to those in the central marrow also derive from distal regions of the BM (i.e. cortical bone, epiphysis or periosteum) (Figure 4f).

Together, these data validate the ability of our approach to identify novel cell types, such as Adipo- and Osteo-CAR cells, and spatially allocate in the BM. Moreover, our data demonstrates that Cxcl12 is mainly synthesized at sinusoidal surfaces by Adipo-CAR cells, but also by Osteo-CAR cells at arteriolar surfaces, and in some instances in non-vascular regions (see also Figure 6g).

### Spatial relationships of BM resident cell types can be accurately predicted based on scRNAseq data and cell adhesion molecule expression

How spatial relationships of cell types are established and maintained within complex organs, such as the bone marrow, remains poorly understood. It has been suggested that the expression of cell adhesion molecules represents an important mechanism that translates basic genetic information into complex three-dimensional patterns of cells within tissues^35^, but this hypothesis has never been investigated systematically in complex systems, due to a lack of spatial and molecular tissue maps. To investigate whether cell type-specific localization of BM populations can be predicted by the differential expression of cell adhesion molecules, we compiled a comprehensive list of well-annotated cellular adhesion receptors and their cognate plasma membrane or ECM-bound ligands and developed the RNA-Magnet algorithm (Figure S10a, Table S3, Methods). RNA-Magnet predicts potential physical interactions between single cells and selected ‘attractor’ populations, based on expression patterns of cell surface receptors and their cognate surface expressed binding partners. For each single cell, RNA-Magnet provides scores for the strength of attraction (RNA-Magnet adhesiveness) and a direction, indicating the attractor population the cell is most attracted to (RNA-Magnet location). To investigate whether RNA-Magnet is capable of recapitulating the spatial relationships of BM cell types, we introduced four anchor populations representing the following niches: osteoblasts for the endosteal niche, sinusoidal endothelial cells for the sinusoidal niche, as well as arterial endothelial and smooth muscle cells both representing arteriolar niches (Figure 5a). Predicted adhesiveness of BM populations to distinct niches strongly correlated with their degree of differential localization as measured by spatial transcriptomics (Figure 5b). Strikingly, cell type-specific localization was also recapitulated with high accuracy for almost all populations (Figures 5c, S10b-d), including the differential localization of Adipo-CAR and Osteo-CAR cells to sinusoidal and arteriolar endothelia, respectively. Interestingly, smooth muscle cells were most attracted to arterial endothelial cells, whereas arteriolar fibroblasts adhered to smooth muscles, recapitulating the consecutive layering observed in blood vessels with the tunica intima (endothelial cells) surrounded by the tunica media (smooth muscle cells), and the tunica externa (ECM produced by arteriolar fibroblasts)^36^. Together, these observations demonstrate the ability of RNA-Magnet to predict spatial localization from single-cell gene expression data and highlight the importance of cell adhesion proteins for tissue organization in the BM.

**Figure 5.**
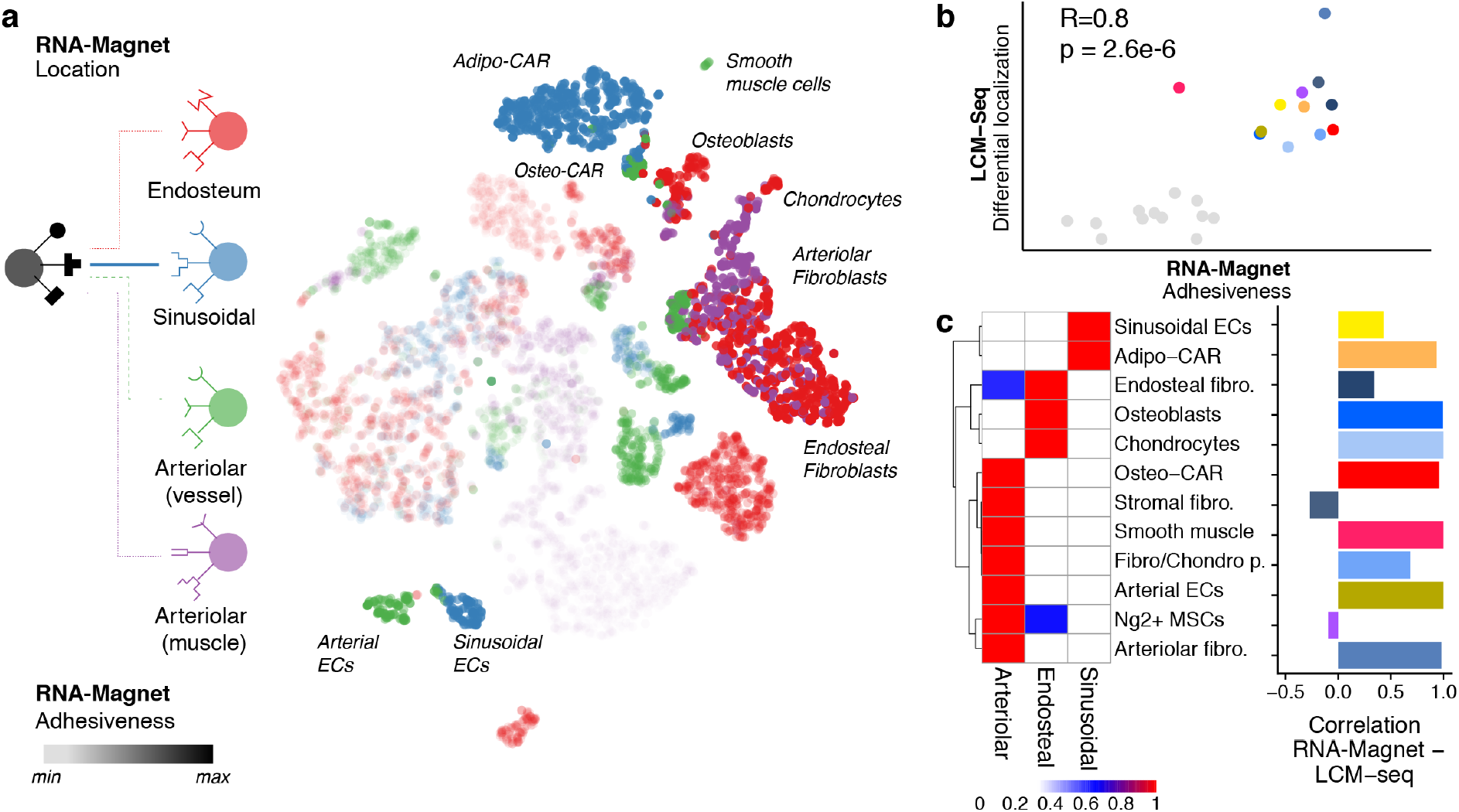
Inference of cellular interactions from single-cell gene expression data by RNA-Magnet. **a**, t-SNE highlighting the cell type each single cell is most likely to physically interact with (RNA-Magnet: Location, indicated by colour), and the estimated strength of adhesion (RNA-Magnet: Adhesiveness, indicated by opacity). **b**, Scatter plot comparing the estimated strength of adhesion (RNA-Magnet score) to the degree to which each cell type is differentially localised between niches (spatial transcriptomics, see also Figure 3c). **c**, Left panel: Heatmap depicting a summary of inferred localisation based on RNA-Magnet. Fraction of cells assigned to a certain niche is colour-coded. Right panel: Bar chart indicating the correlation between the RNA-Magnet estimate of localization and the LCM-seq estimate of localization.

### Cellular and spatial sources of cytokines and growth factors in the BM

Key biological processes occurring in the bone marrow are thought to be mediated by the coordinated action of a diverse set of cytokines and growth factors. However, the identity of cytokine-producing cells and their organization into spatial and functional BM niches remain poorly understood. In particular, the cellular and spatial sources of haematopoietic stem cell (HSC) maintenance factors, such as *Cxcl12* and Scf (*Kitl*), remain controversial ^6–9^ with Lepr-Cre, NG2-Cre, CAR cells, Nestin-dim, osteoblasts and endothelial cells each being separately considered as potential main sources of these factors. To identify cells serving as the source of HSC maintenance factors, we initially examined our scRNAseq dataset. Our data show that *Cxcl12* and Scf are indeed expressed by arterial endothelial ^37^ and some mesenchymal cell types^9^. However, their expression was several orders of magnitude higher in the Adipo- and Osteo-CAR populations (Figure 6a, see also Figure 4). In order to confirm Cxcl12 expression from Adipo- and Osteo-CAR cells at protein level, we developed FACS marker strategies to discriminate these cells types, and confirmed them by FACS-based index single-cell RNAsequencing (Figure 6b). Comparative analyses demonstrated that Adipo-CAR cells are CD45^neg^CD71^neg^Ter119^neg^CD41^neg^CD51^+^VCAM1^+^CD200^mid^CD61^low^, whereas Osteo-CARs and NG2^+^ MSCs expressed CD200 and CD61 at high levels (Figure 6c, S11). Intracellular flow cytometric analyses confirmed that both CAR populations are the main producers of Cxcl12 protein, whereas endothelial cells produced detectable but significantly lower levels (Figure 6d, see also Figure 4a-c). CAR cell populations were also among the main producer of key cytokines required for B cell and myeloid lineage commitment, such as *Il7* and *m-CSF* (*Csf1*) (Figure 6a). Intriguingly, among all BM cell types, CAR cell populations produced the highest numbers of distinct cytokines and growth factors, and attributed the highest proportion of transcriptional activity to cytokine production, suggesting that they act as ‘professional cytokine-producing cells’ (Figure 6e,f). Together, these observations suggest a model in which differential localization of professional cytokine producing cells to cellular scaffolds results in the establishment of specific micro-niches. In line with this, spatial transcriptomics revealed that the five niches investigated showed unique production patterns of cytokines and growth factors (Figure 6g,h). Importantly, CAR cell-derived cytokines were predominantly produced in arteriolar and sinusoidal niches, and net production of growth factors and cytokines was significantly higher in vascular if compared to non-vascular niches, in line with the preferential localization of Osteo- and Adipo-CAR cells to their respective endothelial scaffolds (Figure 6g). Together, these observations suggest that specific localization of professional cytokine producing BM resident cells (such as Adipo- and Osteo-CAR cells) results in the establishment of unique niches with both arteriolar and sinusoidal niches being key sites for the production HSC maintenance and differentiation factors.

**Figure 6.**
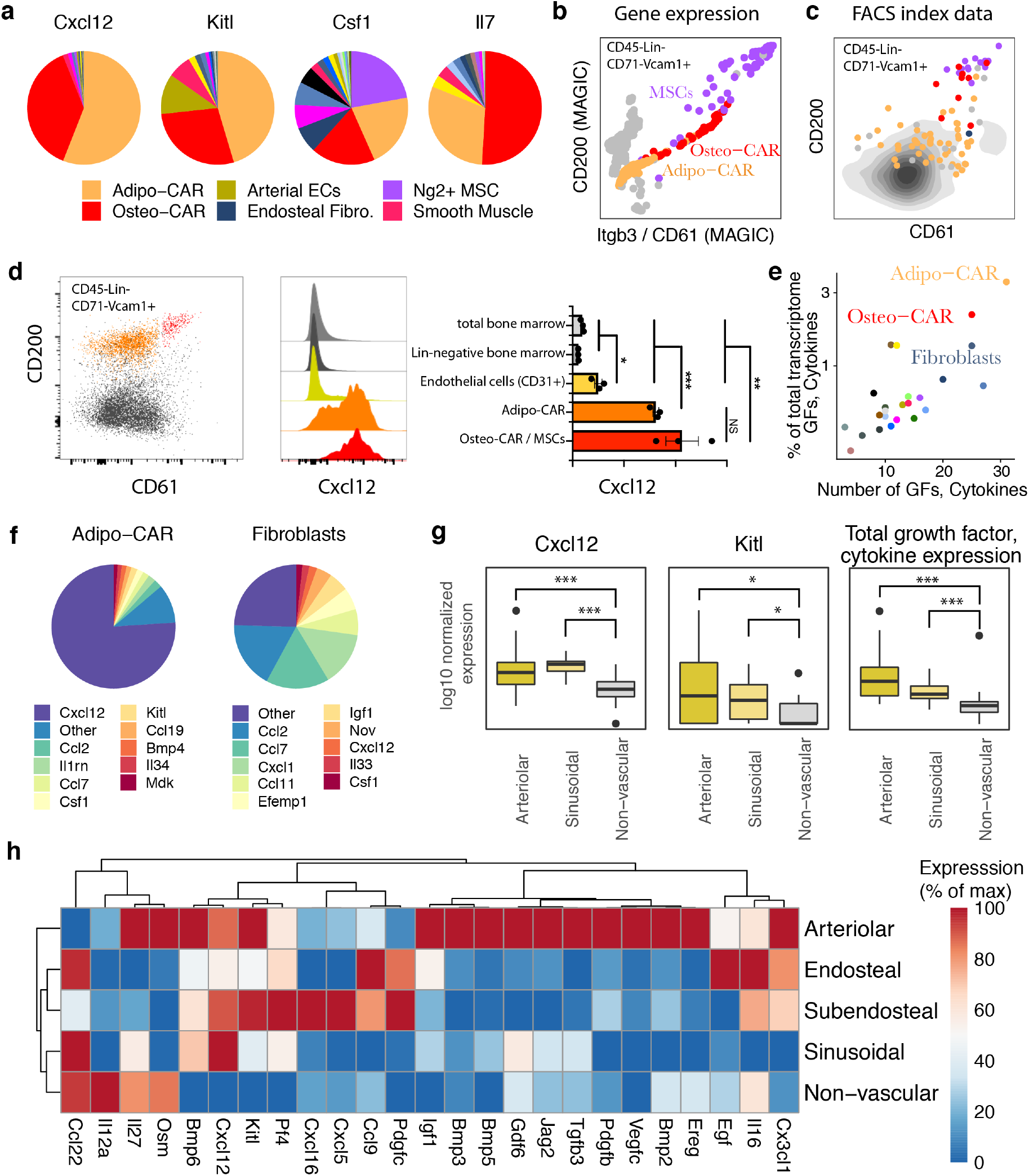
Cellular and spatial sources of key cytokines in the bone marrow. **a**, Contribution of cell types to distinct cytokine pools. Mean gene expression across all cells constituting each cell type is compared. **b**, Single-cell gene expression levels of *Cd200 and Itgb3 (CD61)* from 10x genomics data in CD45^neg^Lin^neg^CD71^neg^Vcam1^+^ cells. MAGIC^47^ was used to impute drop-out values. Relevant populations are colour-coded. **c**, Surface marker levels of CD200 and CD61 from indexed scRNAseq data in CD45^neg^Lin^neg^CD71^neg^Vcam1^+^ cells. FACS index values for n=91 cells subjected to indexed scRNAseq (see methods). The colour indicates the most similar cell type from the main data set as identified by scmap^49^. **d**, Intra-cellular FACS analyses of Cxcl12 expression in total BM, lineage-negative BM, Lin-CD31^+^ endothelial cells and CD61/CD200 subpopulations of CD45^neg^Lin^neg^CD71^neg^CD51^+^Vcam1^+^ CAR cells. Statistics were performed using an unpaired t test. ***: p < 0.001, **: p<0.01*,: p<0.05 **e**, Quantification of the number of growth factors (GFs) and cytokines expressed by each cell type, and the fraction of total mRNA devoted to producing growth factors and cytokines. For a list of growth factors and cytokines used, see Table S3. **f**, Relative expression of cytokines and growth factors in Adipo-CAR cells and Fibroblasts. **g**, Expression of Cxcl12, Kitl, and summed expression of all cytokines and growth factors in arteriolar, sinusoidal and non-vascular niches from spatial transcriptomics data. P-values for differential expression relative to non-vascular niches are from limma/voom^50^ (Cxcl12, Kitl) or a Wilcoxon ranksum test (sum); ***: p < 0.001, *: p<0.05. **h**, Expression of cytokines, chemokines and growth factors in the different niches measured by LCM-seq. Only factors with significant differences between niches are included.

### Systems-level analysis of intercellular signaling interactions of BM resident cell types

To gain a systems-level overview of potential intercellular signaling interactions, we applied RNA-Magnet to soluble signaling mediators (e.g. cytokines, growth factors etc.) and their receptors (Figures 7a, S10e). Unlike previous approaches for the reconstruction of signaling networks from single cell data^38–40^, RNA-Magnet incorporates information on surface receptors with low mRNA expression; is based on a highly curated list of ligand-receptor pairs (Table S3); and specifically identifies the enrichment of signaling interactions between pairs of cell types, with an improved runtime compared to currently available packages^38^ (see also Methods).

**Figure 7.**
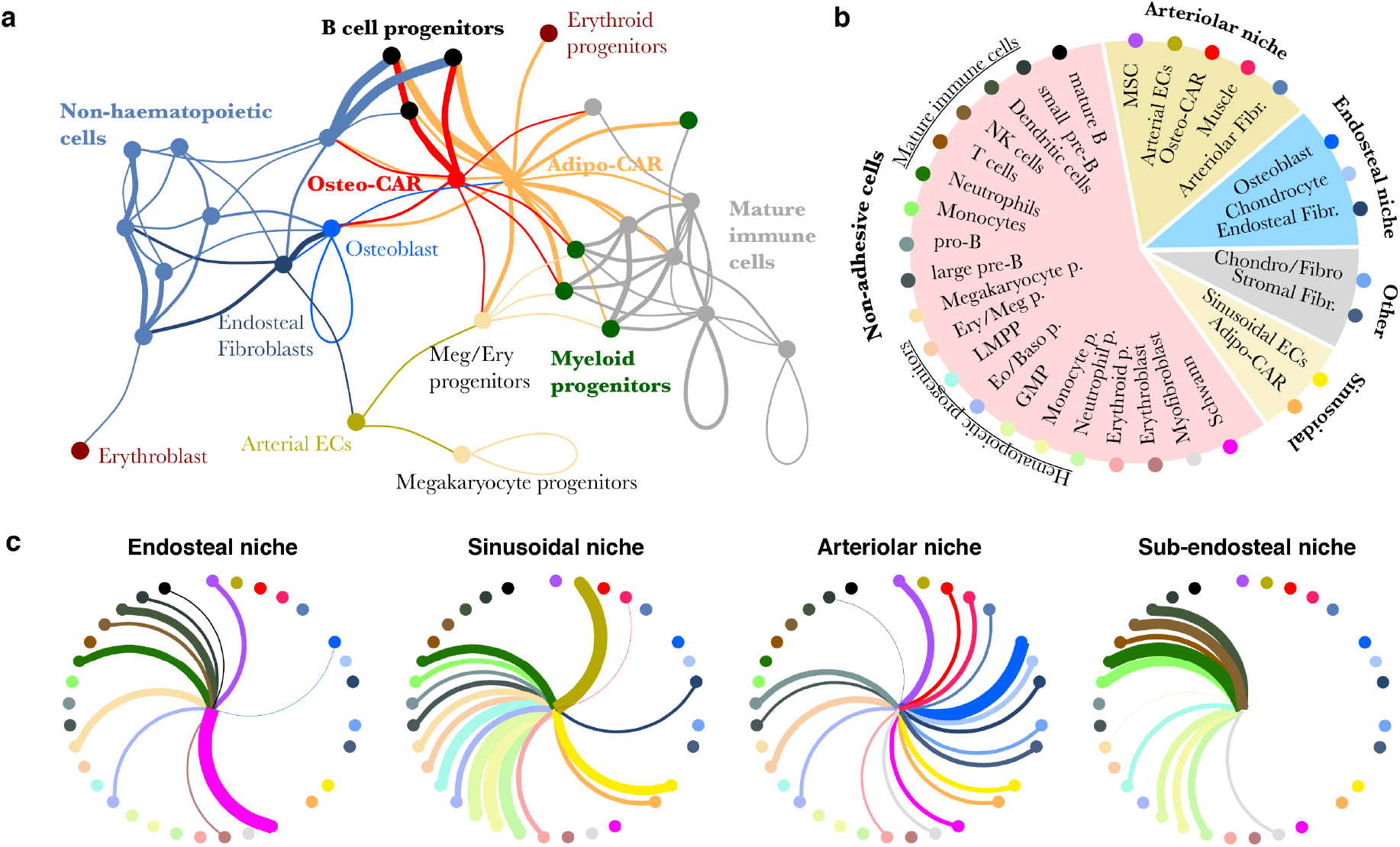
Systems-level analysis of signaling potential in the BM. **a**, Inference of signaling interactions between cell types by RNA-Magnet. If a cell type is enriched in expression of ligands for receptors expressed by a second cell type, a line is drawn between these cell types, with colour indicating the ligand-producing cell type and line width indicating the strength of enrichment. For details, see methods and figure S9e for a fully labelled version of the figure. **b**, Summary of niche composition, as estimated from LCM-seq (see also figure 3f). **c**, Inference of signalling interactions between niches and cell types. Line width indicates the strength of enrichment for expression of ligand-receptors pairs. Cell types are arranged as in subfigure *b*.

The network obtained from RNA-Magnet analyses formed two disconnected signaling clusters consisting of either mature immune or non-haematopoietic cells (i.e. endothelial, mesenchymal and neuronal cells), suggesting that immune and non-immune cells preferentially communicate within their respective groups. For example, many signals potentially sensed by osteoblasts^41^ (e.g. Bmp-, PTHrP-, FGF-signalling, see Figure S10f) are released by the newly identified endosteal fibroblasts described above. In contrast to mature immune cells, HSPC populations frequently received signals from non-haematopoietic cells, suggesting that HSPCs gradually switch from a mesenchymal to an immune signaling niche upon lineage commitment. Importantly, both CAR cell populations were the most important source of signals sensed by all myeloid and lymphoid progenitors (Figure 7a). In accordance with the specific localization of Osteo- and Adipo-CAR populations to arteriolar or sinusoidal scaffolds, an analysis of the net signaling output of distinct local niches implied that lymphoid and myeloid progenitors receive strong input from cytokines produced in vascular and especially sinusoidal niches (Figure 7b, c). Together, these analyses support a concept where distinct biological BM processes are mediated by specific combinatorial signaling input from different local niches^42,43^ and provide a systems-level view of signaling interactions in the BM.

## DISCUSSION

In this study we have combined single-cell and spatially resolved transcriptomics in order to transcriptionally map all major BM resident cell types, identify novel cell types and spatially allocate them to distinct BM niches. Our data clarifies the cellular and spatial origin of key cytokines regulating BM haematopoiesis. For example, the key HSC factors Cxcl12 and Scf are mainly produced by two newly described subpopulations of previously known CAR cells, which we have termed Osteo- and Adipo-CAR cells, according to their gene expression profile. Besides stem cell maintenance factors, CAR cell subsets produce the highest amounts of cytokines among all BM resident cell types, including main cytokines mediating myeloid and lymphoid differentiation, in line with a recent study demonstrating that IL7 and CXCL12 are produced by the same BM cell type^44^. This suggests that CAR cells act as ‘professional cytokine producing cells’ and constitute central niche cells orchestrating many aspects of haematopoiesis. While the more abundant Adipo-CAR cells localize to sinusoidal endothelia, Osteo-CAR cells localize to non-vascular regions or extensively cover arteriolar endothelia. Accordingly, high production of stem cell maintenance and differentiation factors could be observed in both sinusoidal and arteriolar niches if compared to non-vascular niches. Together, this suggests that vascular (both arteriolar and sinusoidal) scaffolds represent the key sites for the production of factors required for stem cell maintenance and differentiation, with the Adipo- and Osteo-CAR cell subsets constituting central cellular hubs.

Distinct cell populations have been suggested to act as mesenchymal stem cells (also known as skeletal stem cells), including Ng2^+^, Nes^+^, LepR^+^ and CD45^neg^CD51^+^CD200^+^ cells^7,10,45,46^. While these populations all constitute heterogeneous populations, pseudo-time analyses of our data suggest that a population of Ng2- and Nes-expressing MSCs resides at the apex of distinct mesenchymal lineages. Importantly, index scRNAseq (Figure 6c, S10b) and comparison to previously published data sets (Figure S6c,d) demonstrated that these cells express most previously described MSC markers (CD45^neg^CD51^+^CD200^+^Ng2^+^Nes^+^Lepr^mid^) and therefore likely represent the true mesenchymal stem cell.

Two recent studies performed single-cell RNA-seq of mesenchymal cells from mouse BM^21,27^. In particular, ref.^27^ complements the scRNA-seq data presented here, since cells from flushed BM were sorted using genetic markers (LepR-Cre, VE-Cad-Cre and Col2.3-Cre). A comparison with our data set reveals that both CAR cell populations as well as MSCs are contained within the LepR^+^ cell compartment (Figure S6). Conversely, our dataset is more comprehensive due to the inclusion of all non-hematopoietic, as well as hematopoietic stem and progenitor cells, therefore enabling the analysis of intercellular signaling. Most importantly, the integration of scRNAseq and spatial transcriptomics enables the systematic localization of cell types to BM niches and the *in situ* measurement of cytokine mRNA synthesis.

Conceptually, our data supports a model where the establishment of unique niches is mediated by differential localization of professional cytokine producing cells to cellular scaffolds. Distinct niche milieus might differentially regulate hematopoietic activities, in line with recent data from genetic studies^42^. In the future, it will be of interest to investigate whether such extrinsic, niche-driven variations determine early fate decisions of hematopoietic stem cells^23^. Overall, our approach, which combines single-cell RNAseq and spatially resolved transcriptomics, reveals the molecular, cellular and spatial organization of BM niches and offers a novel and broadly applicable strategy to systemically map the organization of whole organs.

## Acknowledgment

We thank M. Milsom, A. Grozhik, J. Velten, I. Lohmann, B. Velten and members of the Steinmetz, Haas and Trumpp labs for helpful discussions and critically proofreading the manuscript. Cxcl12-GFP mice were originally from T Nagasawa. We thank K. Bauer and J. Mallm from the DKFZ Single-cell open lab, D. Krunic from the DKFZ microscopy core, M. Paulsen from the EMBL flow cytometry core facility, M. Eich from the DKFZ flow cytometry, the EMBL genomics core facility, S. Terjung from the EMBL Advanced Light Microscopy Facility (ALMF), the Carl Zeiss AG, J. Schnell, L. Becker, S. Renders, P. Werner and S. Sood for technical support. This work was supported by the SFB873, FOR2674 and FOR2033 funded by the Deutsche Forschungsgemeinschaft (DFG), the SyTASC consortium (Deutsche Krebshilfe) and the Dietmar Hopp Foundation (all to A.T.), the US National Institutes of Health (P01HG00020527 to L.M.S.), the ERC (294542 to L.M.S) and the José Carreras Foundation for Leukemia Research (DCJLS 20 R/2017 to L.V., A.T., L.M.S. and S.H.).

## Author Contributions

CB and JA developed experimental methods and performed the majority of experiments with conceptual input from SH, LV, AT and LMS. LV analysed the data with conceptual input from SH and the other authors. LV, CB and LMS developed RNA-Magnet. SH and LV conceived the study, supervised experimental work and wrote the manuscript, with contributions from CB, JA, AT and LMS. FG and PHM experimentally supported this work. PMH and CAN performed whole mount imaging. All authors have carefully read the manuscript.

## Supplementary Information

Supplementary tables S1-S3 are provided as xls files. Supplementary figures S1–S11 and supplementary table S4 are provided below in tables and figures, respectively. A supplementary note is contained at the end of this document.

---

**Figure S1.**
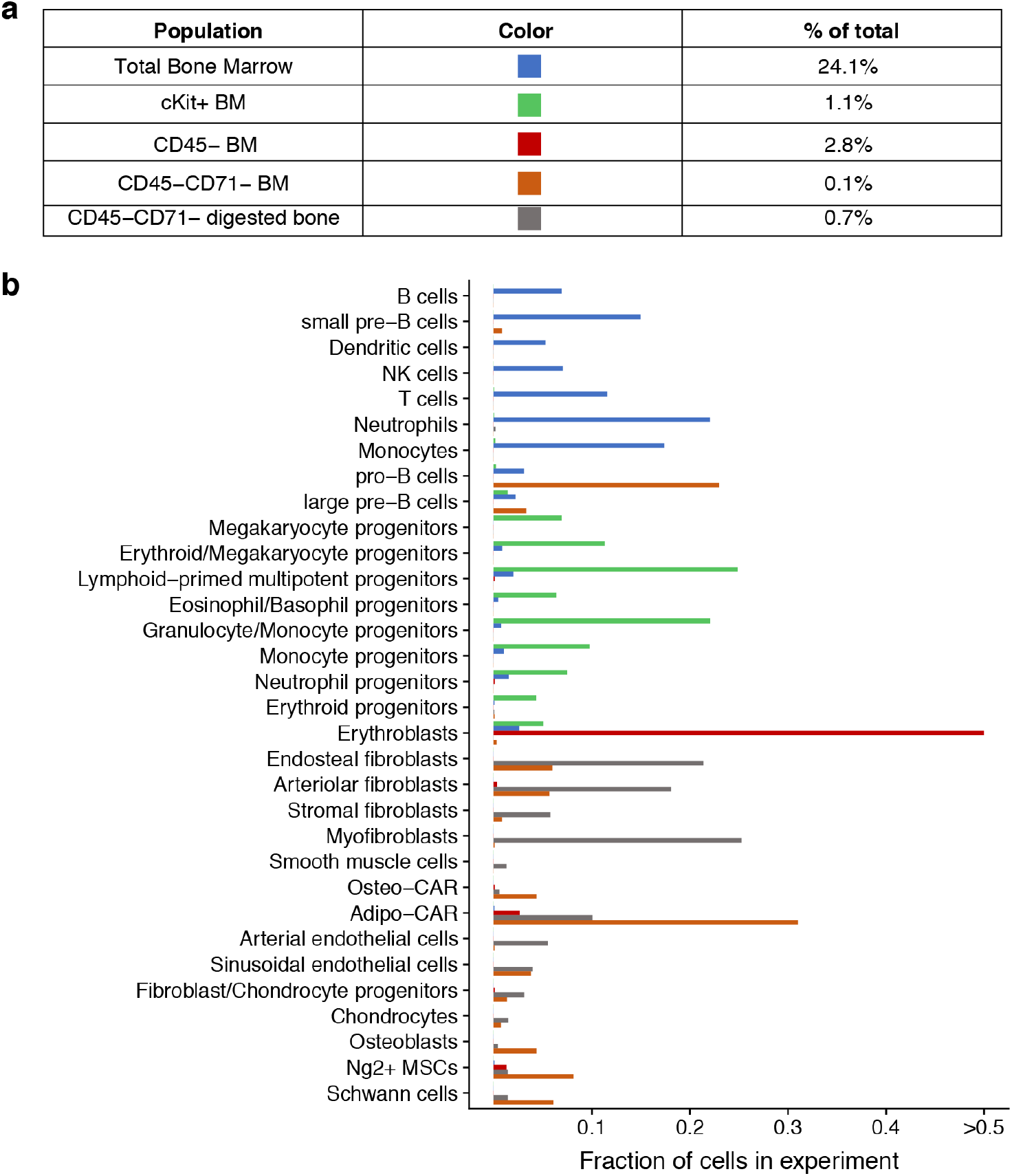
Cellular composition of populations defined by flow cytometry. **a**, Abundance of different gates as fraction of total. **b,** Quantification of cell type composition for each FACS gate shown in main figure 1a.

**Figure S2.**
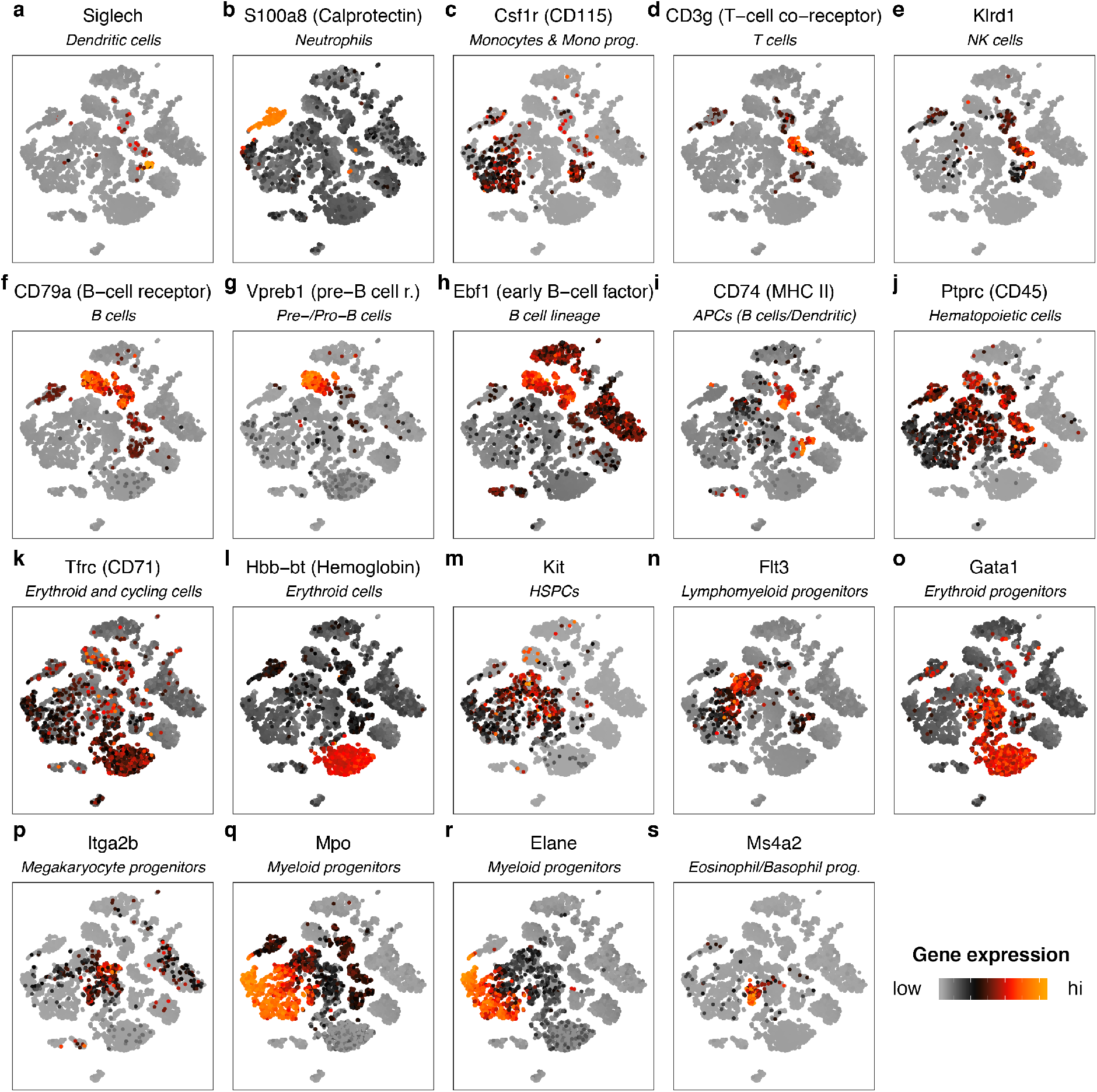
Expression of marker genes for haematopoietic populations highlighted on t-SNE. For full lists of marker genes, see table S1.

**Figure S3.**
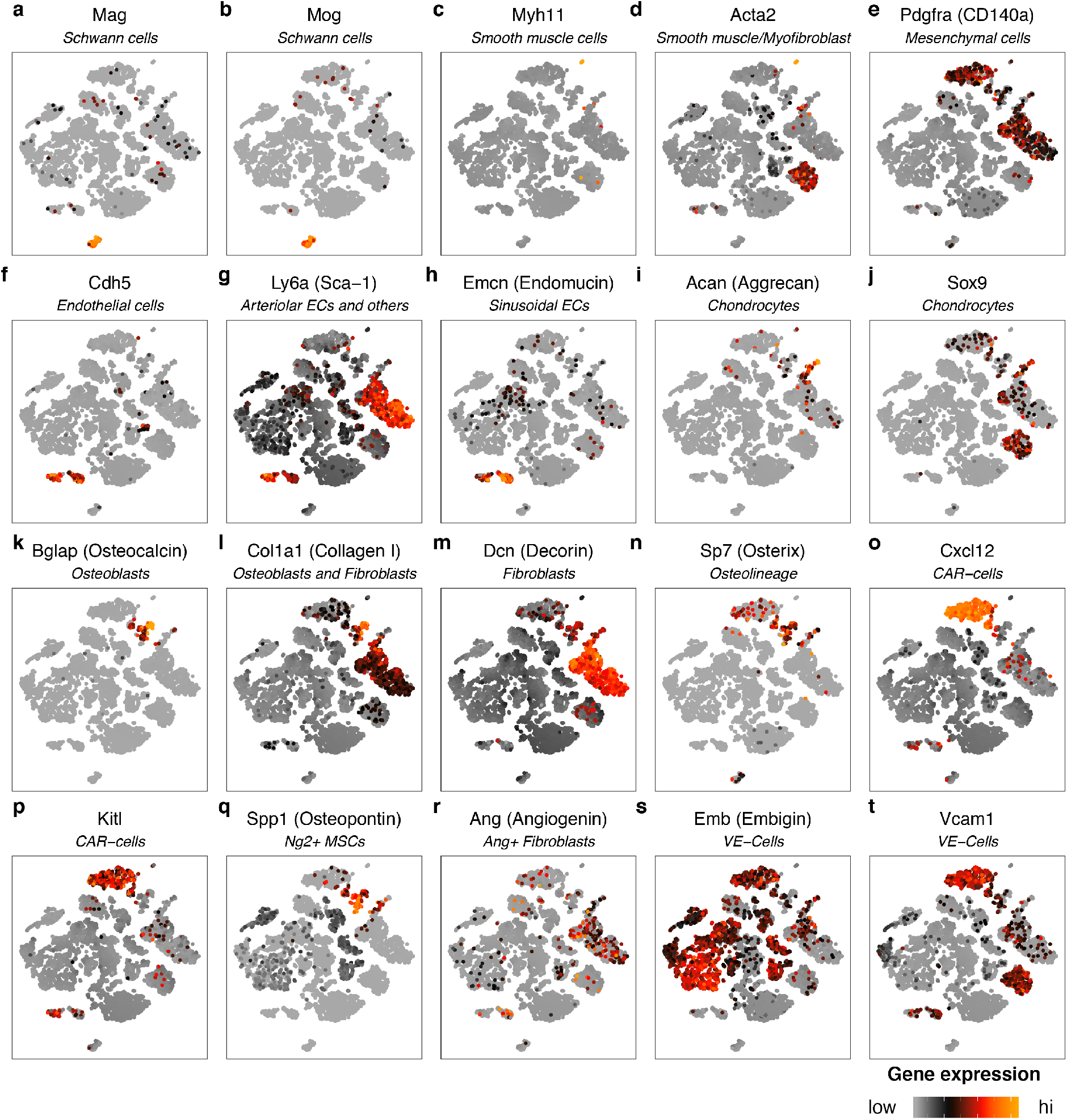
Expression of marker genes for non-haematopoietic populations highlighted on t-SNE. For full lists of marker genes, see table S1.

**Figure S4.**
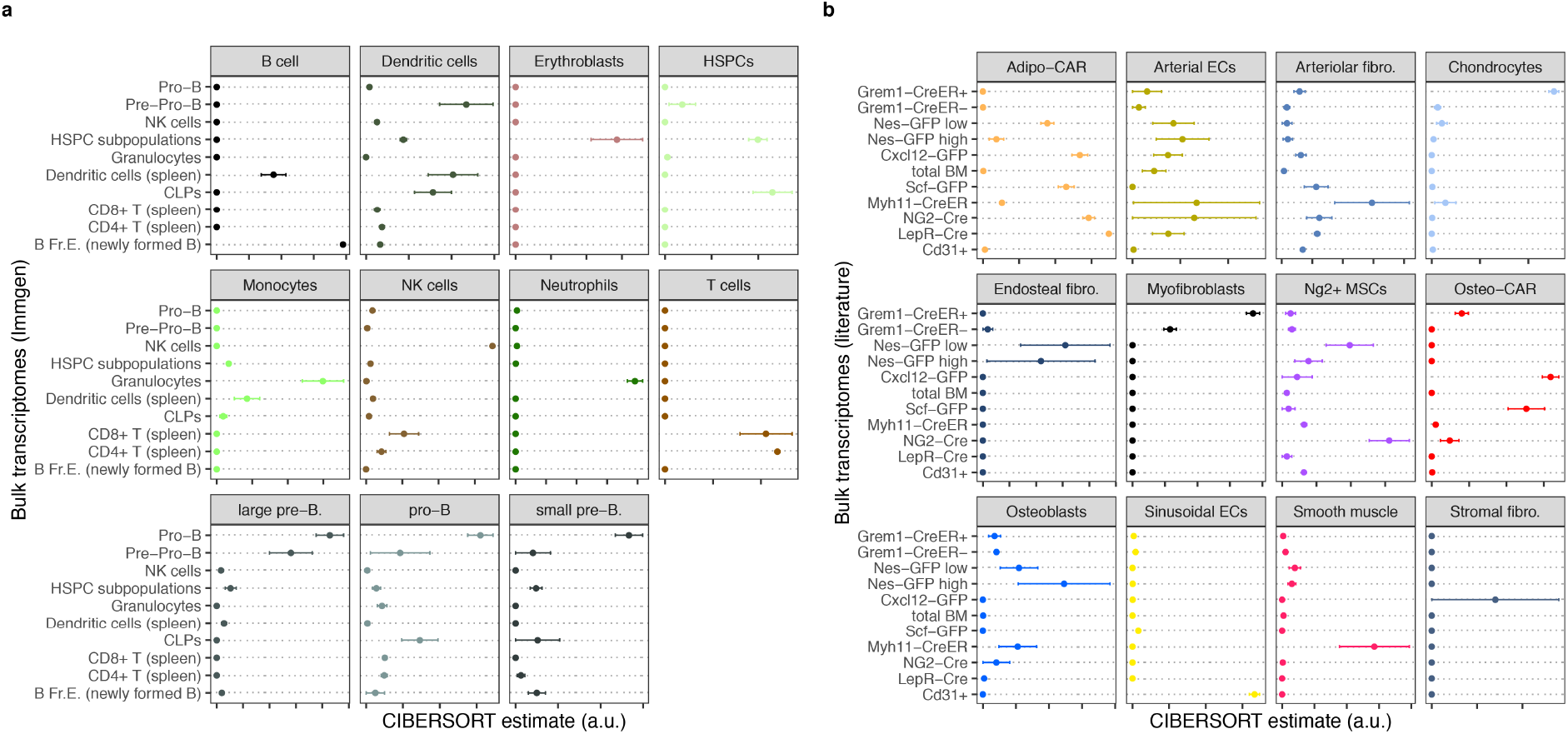
Comparison of cell type transcriptomes determined by scRNA-seq to data from bulk populations described in literature. **a**, Enrichment of gene expression signatures of haematopoietic populations in immune cell transcriptomes published by the immgen consortium (data source: GEO GSE109125)^51^. **b**, Enrichment of gene expression signatures of non-haematopoietic populations in published transcriptomes of populations defined by genetic markers^5,7–9,52^; see methods for specification of data sources, and see the supplementary note for a detailed evaluation of the algorithm used. Error bars indicate standard error of the mean for n=3 to n=6 bulk transcriptome samples per class.

**Figure S5.**
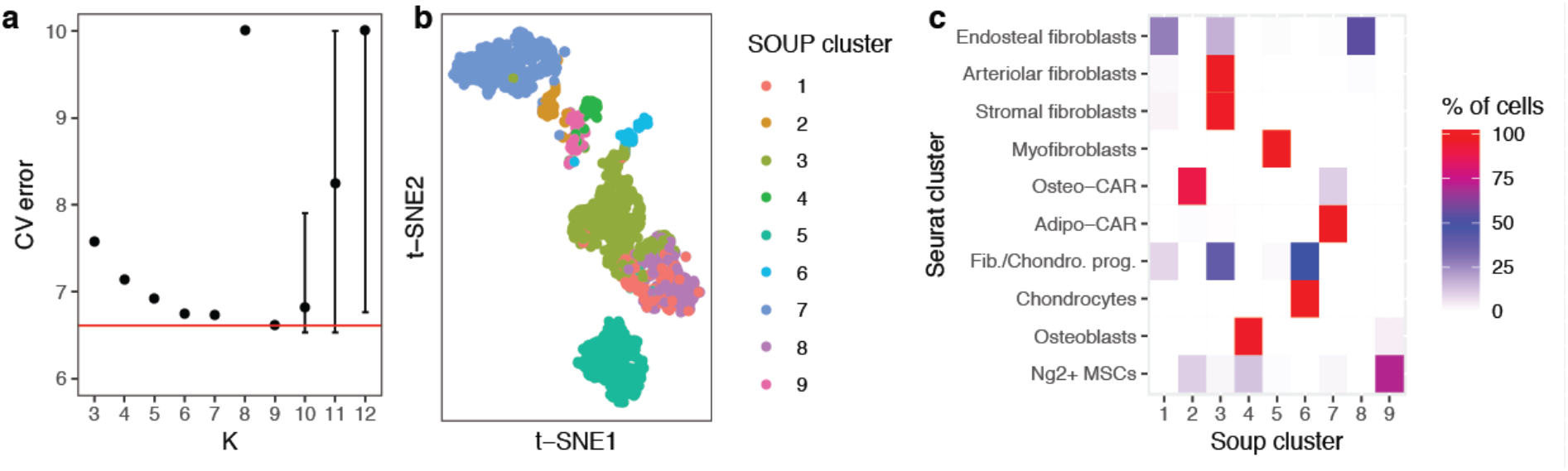
Comparison of clustering methods. **a**, The optimal number of mesenchymal cell clusters was determined using the SOUP method^15^, a semi-soft clustering algorithm designed to distinguish between distinct cell types and transition states between cell types. **b**, Main cluster identity from SOUP highlighted on the t-SNE from figure 1b (mesenchymal cell types only). **c**, Comparison of clusters identified by Seurat (Figure 1b) to clusters identified by SOUP (Figure S5b) demonstrates strong overlap between both methods.

**Figure S6.**
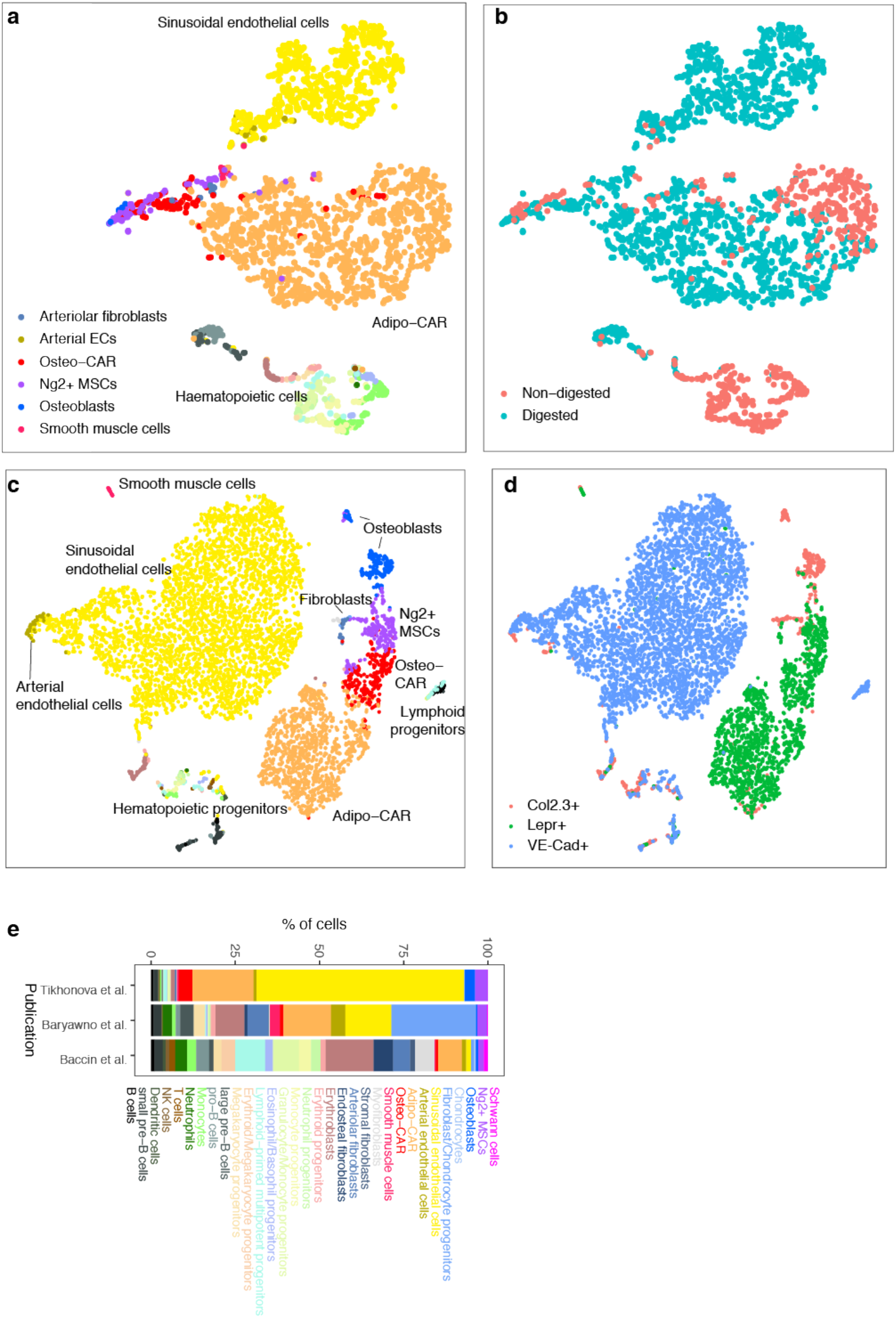
Comparison of cell isolation methods and reference datasets. **a**,**b**, Additional single-cell RNA-seq data was generated as described, except that bone marrow was derived by flushing bones and subjected or not subjected to enzymatic digestion. Data was projected to two dimensions using t-SNE and cell type labels were assigned using the anchoring approach implemented in seurat3^53^. **c**,**d**, Single-cell RNA-seq data from a recent study of different genetically labelled populations from flushed bone marrow^27^ was projected to two dimensions using t-SNE and cell type labels were assigned using the anchoring approach implemented in seurat3^53^. **e**, Comparison of cell type frequencies between various published datasets.

**Figure S7.**
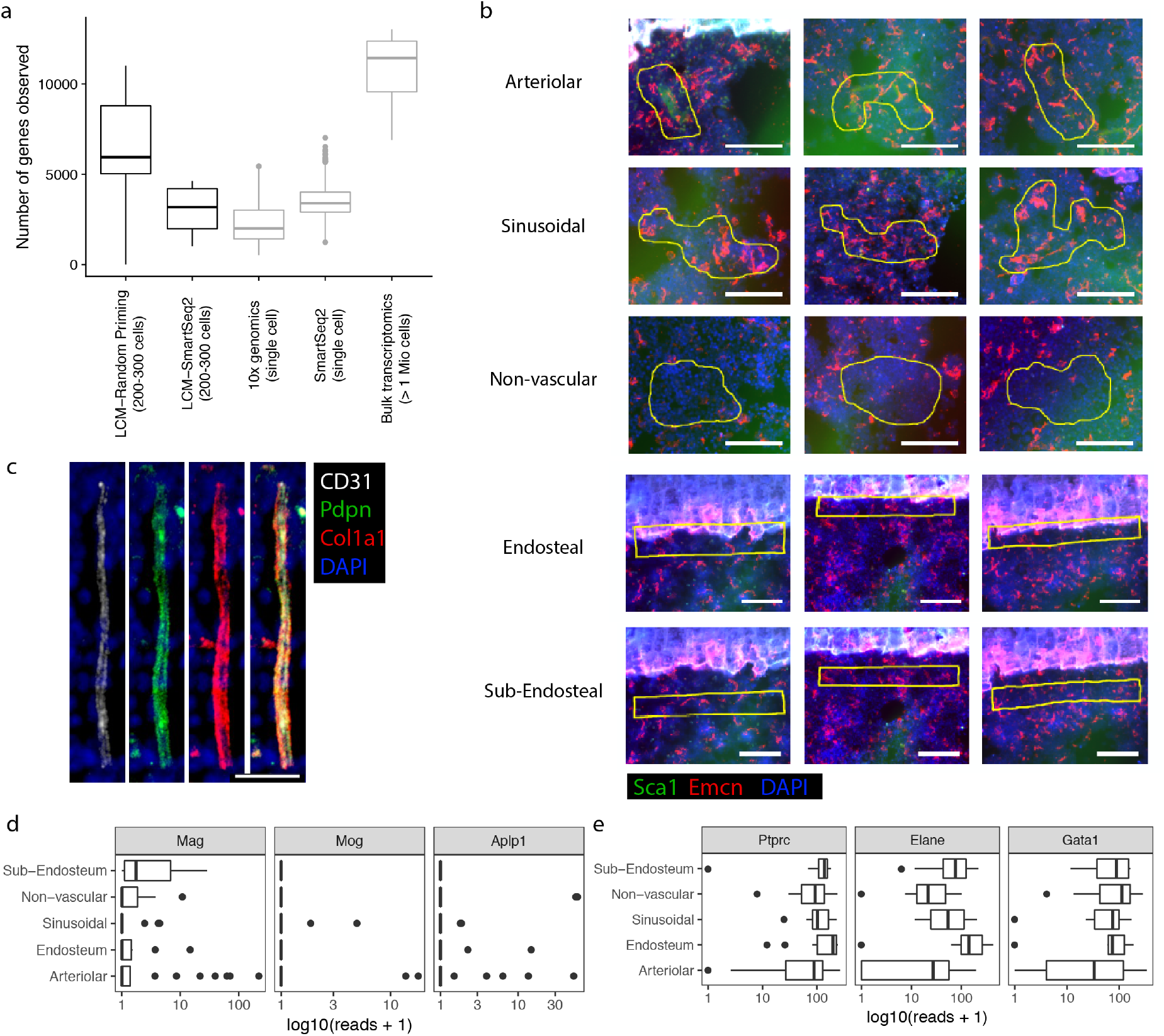
Technical properties of the LCM-seq dataset. **a,** Boxplots comparing the number of genes observed per sample in different protocols. All samples were down-sampled to 1 million reads for comparison. For the dataset presented in main figure 3, the protocol relying on random priming was used. **b**, Representative images of samples collected for LCM-seq; scale bar corresponds to 100 μm. **c**, Immunofluorescence staining of a BM arteriole stained for Colla1, Pdpn and CD31. Scale bar: 20 μm. **d,** Schwann cell markers were lowly expressed across all niches **e,** haematopoietic markers were highly expressed across all niches.

**Figure S8.**
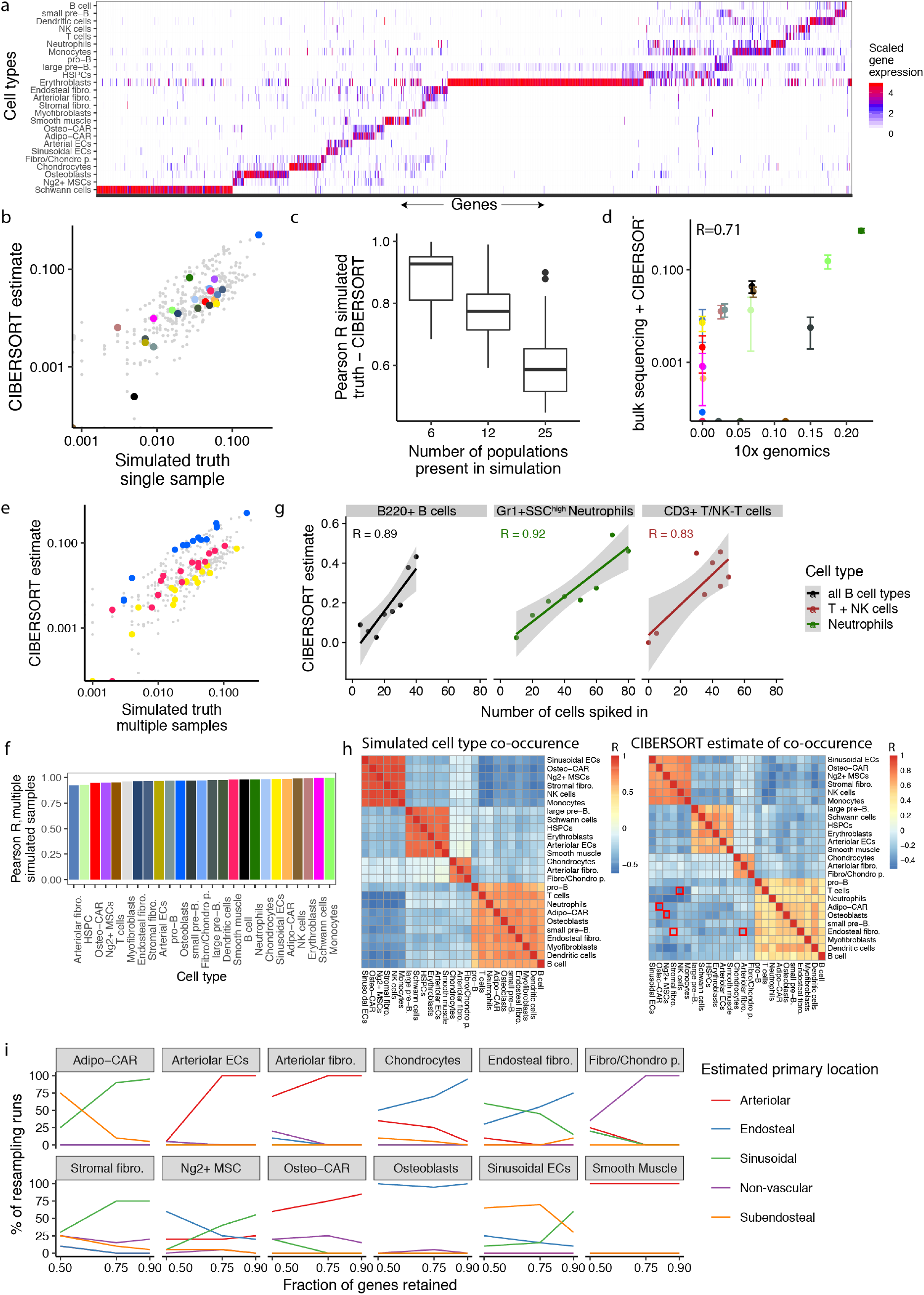
Evaluation of the CIBERSORT algorithm, see also supplementary note. **a**, Heatmap of population-specific marker genes used for the algorithm. **b**,**c**, Simulations to assess the ability of CIBERSORT to decompose individual samples; see supplementary note for detail. **d**, CIBERSORT estimates of cell type composition of total bone marrow, compared to the cell type composition estimate from 10x genomics (see figure S1b). **e**,**f**, Simulations to assess the ability of CIBERSORT to identify changes in population frequencies across multiple samples; see supplementary note for detail. **g**, FACS was used to assemble 8 different pools of B220^+^ B-cells, CD3^+^ T/NK-T cells and Gr1^+^SSC^high^ neutrophils. Each pool contained a total of 100 cells at predefined ratios of B cells, T cells and neutrophils. Pools were then fixed and processed using the LCM-seq protocol, and CIBERSORT was used to decompose their composition. Estimates for T and NK cells, as well as different B-cell subpopulations, were summed for the display. **h**, Simulations to assess the ability of CIBERSORT to discriminate between similar cell types; see supplementary note for detail. Red squares highlight pairs of similar cell types. **i**, Stability of the CIBERSORT estimates from main figure 3e with regard to re-sampling of the marker gene lists used; see supplementary note for detail.

**Figure S9.**
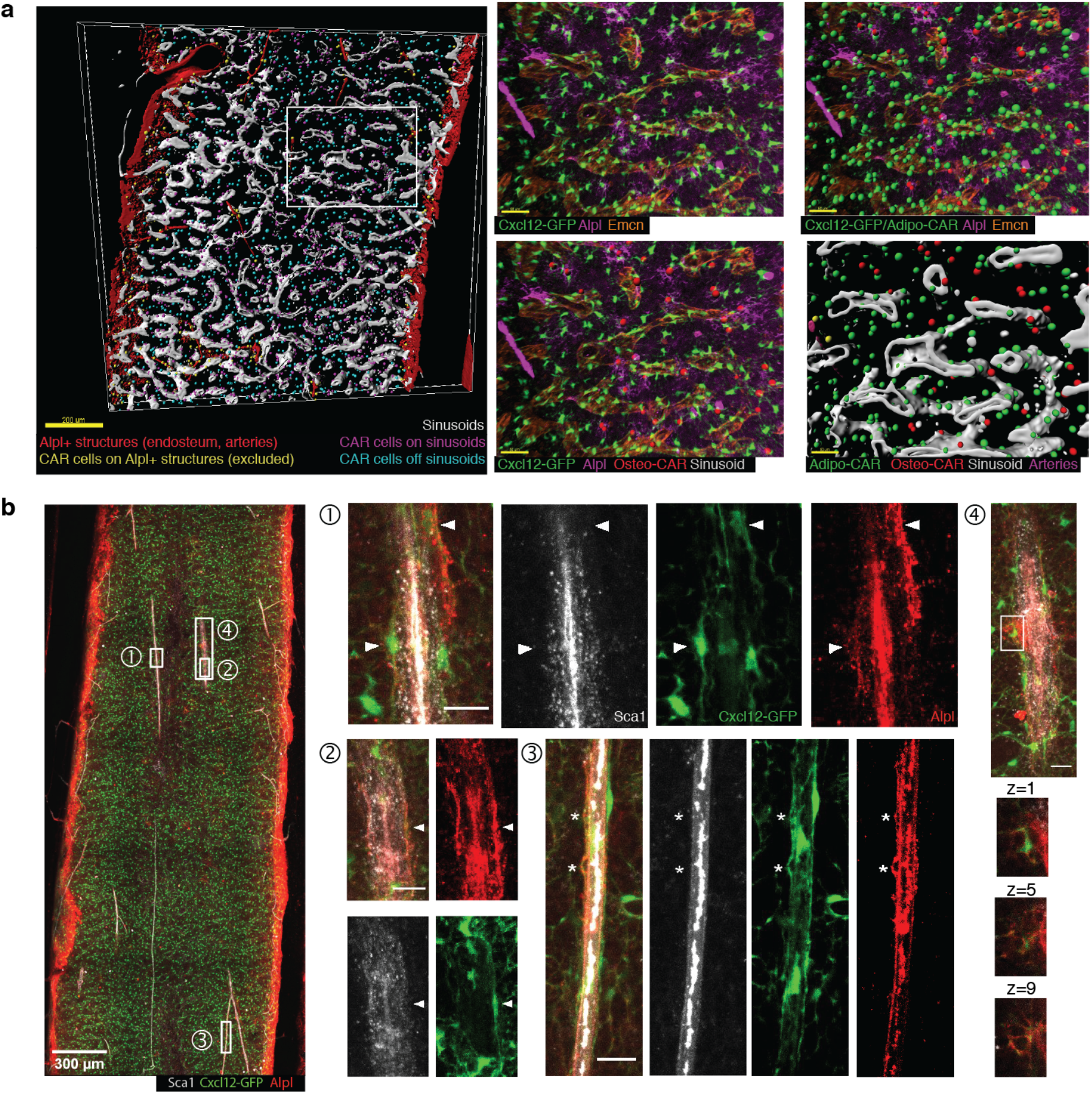
Whole-mount imaging and data analysis. **a**, Whole-mount imaging data of a Cxcl12-GFP bone section stained for Alpl and Emcn was segmented in 3D using the imaris software. Large Alpl^+^ surfaces (red, corresponding to endosteum and arteries) were identified and any GFP^+^ spots with <20μm proximity to these structures were excluded from further analysis (yellow spots). Remaining GFP^+^ spots were classified as within 15μm of sinusoidal vessels (purple dots), of away from sinusoidal vessels (cyan dots). GFP^+^ spots were further classified as Alpl^+^ (right panels, red spots) or Alpl^−^ (right panels, green spots). **b,** Like in main figure 4c. In ROI 3, asterisk correspond to GFP^+^Alpl^+^ protrusions on, but clearly distinct from, Sca1^+^ arteriolar endothelial cells. In ROI 4, various z-sections of a highly reticulate Cxcl12-GFP^+^Alpl^+^ cell are shown.

**Figure S10.**
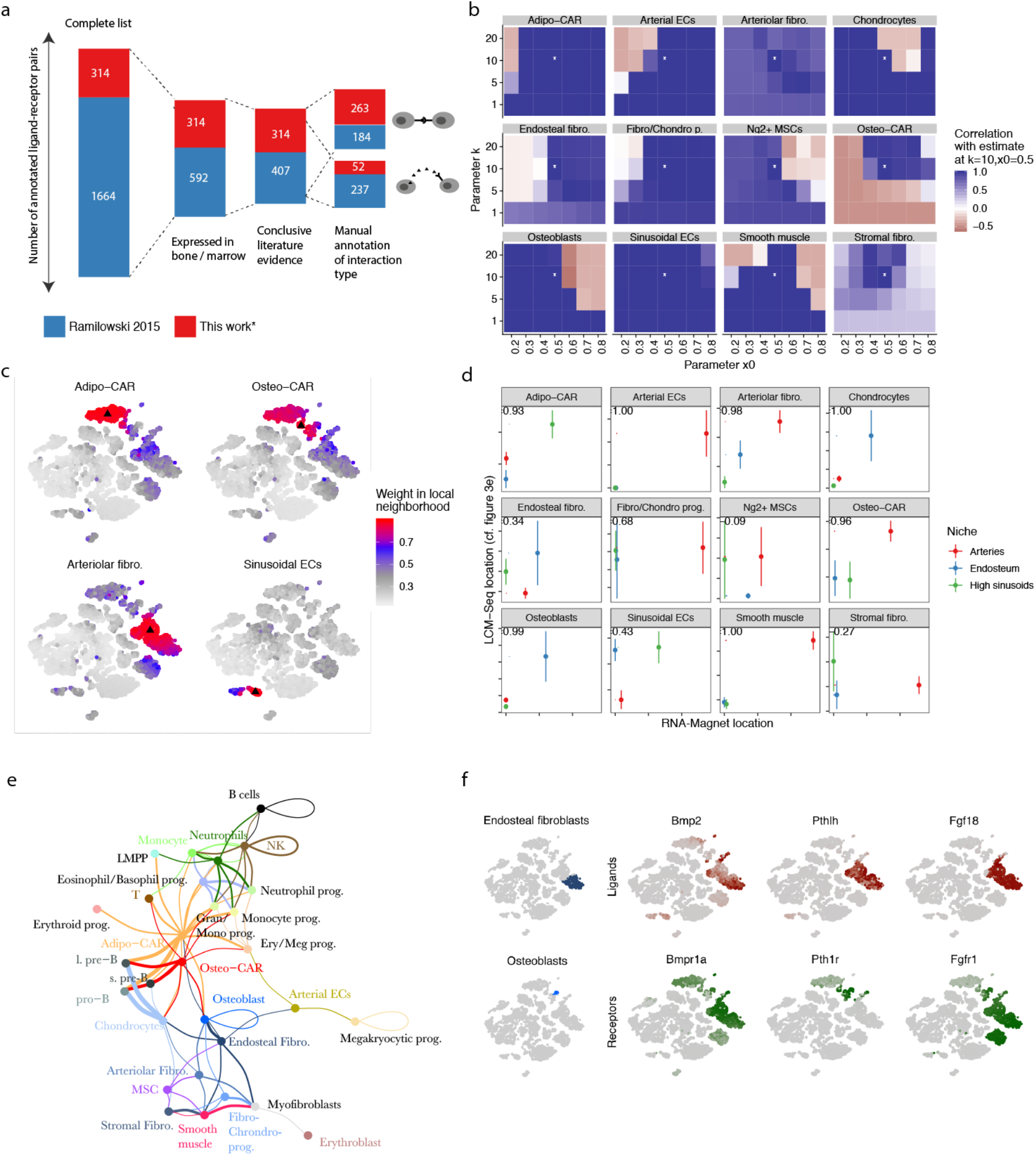
Analyses using RNA-Magnet. **a**, Overview of the receptor-ligand list used. See methods for data sources, and table S3 for the complete list. **b**, Stability of the RNA-Magnet location estimate for different choices of the fuzzification parameters k and x_0_. For each parameter set, RNA-Magnet location estimates were summarised per cell type, and compared to the summarised location estimate displayed in figure 5c. The asterisk indicates the parameter set used in figure 5c. **c**, Choice of local neighbourhoods. As detailed in the methods section, RNA-Magnet works by identifying interactions specific to a single cell compared to similar cells. The figure displays the size of local neighbourhoods for four representative cells demarked by a black triangle. **d**, Detailed comparison of location estimates obtained from LCM-seq and RNA-Magnet. See also main figure 5c. **e**, Fully labelled display of the network from main figure 7a. **f**, Expression of selected cytokines and growth factors involved in bone remodelling.

**Figure S11.**
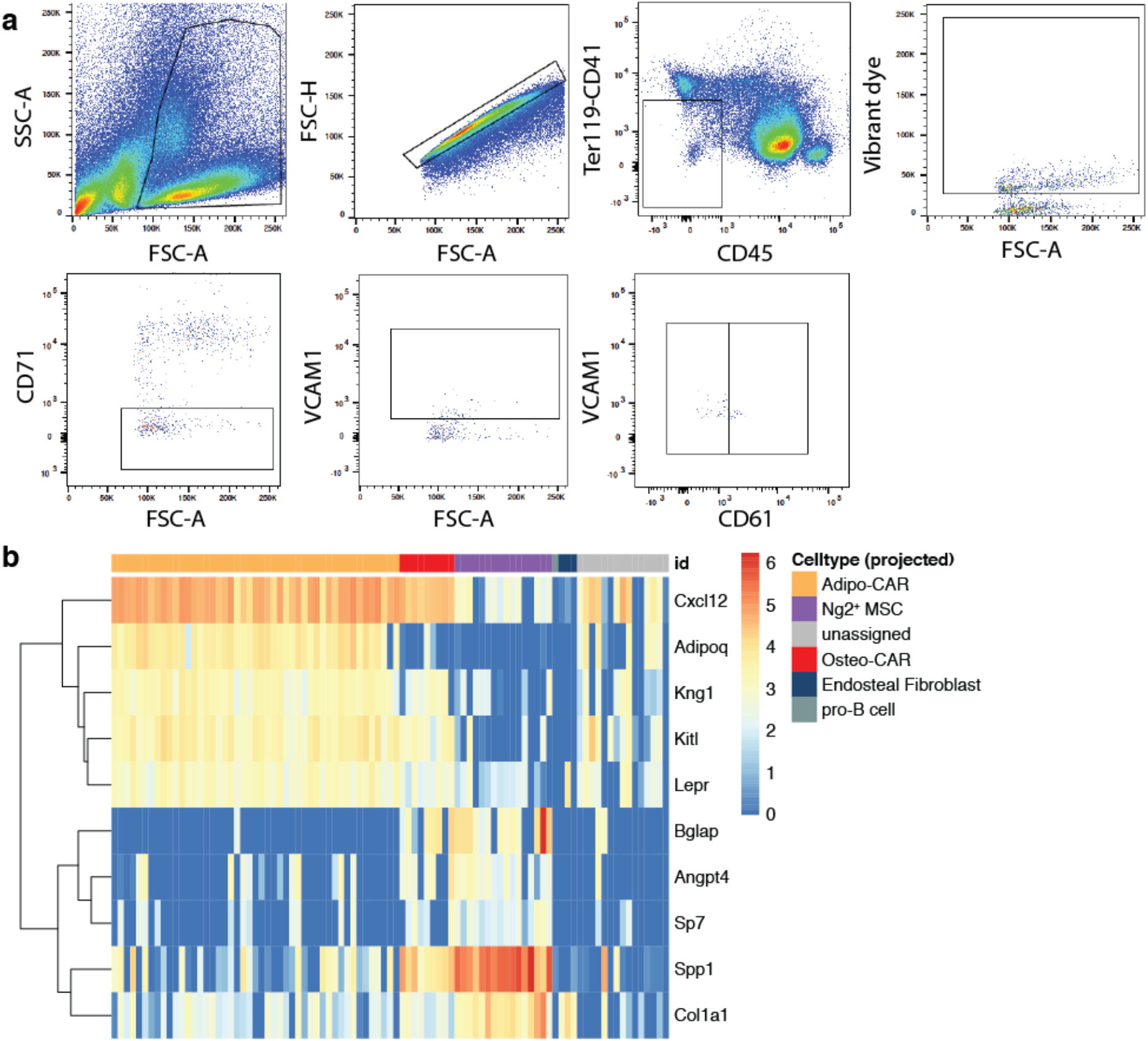
Index-sorting analysis of Lin^neg^Vcam1^+^ cells. **a**, Sorting scheme used **b**, Expression of key marker genes confirm the cell type assignment obtained by scmap, cf. main Figure 6c.

## Methods

### Mouse experiments

Mice were purchased from the distributors Janvier and Envigo, and housed under specific pathogen-free conditions at the central animal facility of the German Cancer Research Center. All animals used were 8-12 weeks old C56Bl/6J females. All animal experiments were performed according to protocols approved by the German authorities (Regierungspräsidium Karlsruhe).

### Tissue harvesting and processing

Femurs, tibiae, hips and spines were dissected and cleaned from surrounding tissue. For all cell sorting and flow cytometry analyses, bones were crushed in cell suspension medium (RPMI 1640 (Sigma) containing 2% fetal bovine serum) using a mortar and a pestle. Dissociated cells were filtered through a 40 μm filter, spun down at 1500 rpm for 5 min, then incubated with 5 ml ACK lysis buffer (Thermo Fisher Scientific) for 5 min at room temperature for red blood cell lysis. Neutralization was achieved with 20 ml cell suspension medium. Cells were then lineage-depleted using Dynabeads Untouched Mouse CD4 Cells Kit (Thermo Fisher Scientific) according to manufacturer’s recommendations, using a home-made lineage cocktail.

For cell extraction from bones, crushed bone chips were washed four times each with 10 ml of cell suspension medium, then incubated with 10 ml of digestion medium (1 mg/ml of each Collagenase II and Dispase in HBSS, all from Gibco) for 30 min at 37 °C in a water bath. Cell suspension was then removed and filtered through a 40 μm filter and the digestion reaction was stopped by adding 40 ml of cell suspension medium. From this point on, cells were treated exactly the same as bone marrow cells above.

### Flow Cytometry

Lineage-depleted bone and bone marrow cells obtained following crushing and digestion were stained with FACS antibodies (Table S4) for 30 min on ice, then washed with cell suspension medium. For intracellular CXCL12 staining, cells were fixed/permeabilized using BD Cytofix/Cytoperm (BD) for 30 min on ice, washed with BD Perm/Wash (BD) and incubated overnight with anti-CXCL12 antibody (1:10), then washed with BD Perm/Wash. All flow cytometric analyses were performed using BD Fortessa flow cytometers. Cell sorting was done using BD Aria I, Aria II and Aria Fusion sorters.

### scRNA sequencing – 10x Genomics

For single-cell RNA sequencing using 10x genomics, bone and bone marrow cells were processed as described above. In addition to FACS markers, cells were stained with a DNA dye (Vybrant™ DyeCycle™ Violet, Thermo Fisher Scientific) to exclude debris and ensure that only cells are sorted for droplet-based scRNA-seq. For this purpose, cells were incubated with 2.5 μl/ml Vybrant dye in cell suspension medium for each 3×10^6^ cells at 37 °C for 30 min in a water bath. Following the incubation, cells were placed on ice and for each experiment, a total of 1.0-1.5×10^4^ events were sorted immediately into 15 μl PBS containing 2% FBS. Cell numbers were confirmed using LUNA™ Automated Cell Counter (Logos Biosystems). 33.8 μl of cell suspension were used as input without further dilution or processing, with final concentrations around 100-200 cells/μl. Reverse transcription and library construction were carried out following the Chromium Single Cell 3’ Reagent v2 protocol (10x genomics, Pleasanton, CA) according to manufacturer’s recommendations. Total cDNA synthesis was performed using 14 amplification cycles, with final cDNA yields ranging from approximately 2 ng/ μ l to 10 ng/μ l. 10x genomics sequencing libraries were constructed as described and sequenced on an Illumina Next-Seq500, with read length 26+58 or 26+98.

### FACS-indexed scRNA sequencing

Lineage-depleted bone marrow cells were obtained by crushing and stained with the following antibodies on ice for 30 min: CD41, CD45, CD51, CD61, CD71, CD200, Ter119 and VCAM1 (Table S4). Indexed single-cells were sorted into 5μl of Smart-Seq2 lysis buffer (2μM Oligo-dT30VN primer, 2mM dNTP mix (10mM each, NEB), 1:50 RNAse inhibitor (promega) and 1:125 Triton X-100 10% (Sigma-Aldrich)) and immediately snap frozen in an ethanol and dry ice bath. Plates were kept at −80 °C until processing. cDNA amplification was performed using a modified Smart-Seq2 protocol by adding, after 3 min at 72 °C, 5μl of RT mix containing 1× SMART First Strand Buffer (Clontech), 2 mM dithiothreitol (Clontech), 2 μM template switching oligo (Exiqon), 10 U μl–1 SMARTScribe (Clontech) and 10 U μl–1 RNASin plus (Promega). Transcriptome amplification was performed using 1X KAPA HiFi HS MM and 0.1μM ISPCR primer, with 21 PCR enrichment cycles. Libraries were constructed using in house produced Tn5^54^ at 1:100 dilution and sequenced on an Illumina Next-Seq 500 sequencer, with 75 cycles single end sequencing.

### Bone preparation and sectioning for immunofluorescent staining

Femurs were dissected and cleaned from muscle, and placed immediately in 4% PFA at 4°C for 30 min. Subsequently, femurs were washed three times in 1X PBS before incubating with 15% sucrose for 2 h, followed by 30% sucrose for another 2 hours. All incubations were performed at 4 °C. Femurs were placed in OCT and frozen at −80 °C until sectioning.

For sectioning, a cryotome (ThermoFisher) was used to generate 12 μm sections at −20 to −22 °C, which were then transferred to slides using the CryoJane Tape-Transfer System (Leica Biosystems). Slides were post-fixed with 4% PFA for 1 min to enhance the adherence of sections to the slide, then washed for 2 min by dipping in PBS. Sections were stained with antibodies in blocking buffer (PBS containing 10% goat serum and 0.2% Triton X100, see table S4 for antibodies used) at 4°C overnight, then washed by dipping into PBS for 1 min. Secondary antibody staining was done in blocking buffer at room temperature for 2 h. Sections were imaged using an LSM710 microscope (Zeiss) equipped with a polychromatic META detector (Lasers: 458, 488, 514, 561, 594, and 633nm), and an Olympus FV3000 Confocal laser scanning microscope with 4 GaAsP spectral detectors, FRAP and FRET (Lasers: 405, 445, 488, 514, 561, 594 and 640 nm). Imaris software (v8.41) was used for data analysis and representation.

### Whole mount imaging of immunostained mouse femurs

The employed whole mount staining protocol has been previously described in detail^11^. Briefly, femurs were isolated, the surrounding connective and muscle tissue carefully removed and fixed in 2% paraformaldehyde in PBS (6h, 4°C). After dehydration in 30% sucrose in PBS (72h, 4°C) the femurs were embedded in OCT medium, snap-frozen and bi-sectioned using a cryotome. The remaining thick bone marrow slice was incubated in blocking solution overnight (0.2% Triton X-100, 1 % bovine serum albumin, 10% donkey serum, in PBS) at 4°C and afterwards stained with primary and secondary antibodies (see table S4) diluted in blocking solution for 2-3 days each, including three one hour washing steps with PBS in between. The samples were immersed in RapiClear 1.52 for 6 hours to increase optical transparency. Images were acquired on an SP8 Leica confocal microscope system. Imaris software (v8.41) was used for data analysis and representation.

### Bone sectioning for LCM-seq

Femurs for LCM-seq were carefully processed following guidelines of good RNA work practice. Femurs were harvested and cleaned as quickly as possible, then placed immediately in ice-cold 4% PFA, fixed for 30 min, and dehydrated in 15% and 30% sucrose solutions, prepared using RNase-free sucrose powder (Acros Organics), for 2 h each. All incubation steps were carried out on ice. Femurs were then flash-frozen in a 2-Methylbutane and dry ice bath and stored overnight at −80°C. Bone sectioning for RNA retrieval was performed in a sterilized cryotome, blades were wiped with RNaseZAP (Sigma) and slides were stored on dry ice until staining (on the same day). For staining, we adapted a shortened immunostaining protocol^55^ starting with thawing the slide quickly at room temperature, then incubating with the primary antibody (1:20 to 1:40, depending on the antibody) on an aluminum rack on ice for 10 min, washing in ice-cold PBS for 30 s, then adding the secondary antibody for 5 min on ice. Higher antibody concentrations may be needed for antibodies of lower quality. Final wash and dehydration were done as follows: dipping for 30 s in each ice-cold PBS, RNase-free H2O, 70%, 95% and 100% ethanol, in this order.

### LCM-seq

Bone sections were processed using the Zeiss PALM MicroBeam Axio Observer Z1 (Zeiss), with a monochromatic Axiocam 506 mono camera, shooting laser at 355 nm and filter sets FS18 C adv (Dapi), FS44 C adv (FITC) and FS45 C adv (mCherry). Image acquisition and sample isolation were carried out using a LD Plan-NEOFLUAR 20X/0.4 objective with the adjustment ring set to 1 (Zeiss). Cutting energy was set to 45 and focus to 67, while laser pulse catapult (LPC) energy was used with delta of 12. For cutting and shooting we used the “CloseCut + AutoLPC” option. No more than 4 sections were processed and scanned in parallel, keeping collection time after staining under 30 min. A 3-channel colour image was acquired for each sample, as well as before/after LPC images and metadata including day of collection, slide number and distance of the area of interest from the bone lining. Areas of 14500 μm^2^ on average, corresponding to around 200-300 cells, were isolated from different bone marrow districts and collected separately in 200 μl AdhesiveCap opaque Eppendorf tubes. After LPC, each collection lid was covered with 15μl of a 1:16 dilution of proteinase K in PKD buffer (Qiagen), incubated at room temperature for 5 min and subsequently snap frozen in dry ice and stored at −80°C overnight. For reverse crosslinking, samples were thawed for 5 min at room temperature, spun for 30 s to collect all the liquid at the bottom and incubated for 1h at 56°C on a PCR block^56^. After incubation, samples were resuspended in 100 μl TRI Reagent (Sigma-Aldrich) under a laminar flow hood and stored in 8-PCR-strips at −80°C until RNA extraction.

RNA extraction and library construction were performed in batches of 16. In brief, after thawing and spinning, each sample was transferred to a 1.5 ml Eppendorf tube and 20 μl chloroform were added. Phase separation was achieved at room temperature after vigorous shaking and spinning at 12500 rpm for 5 min, 40 μl of aqueous phase were collected from each sample and added to 75.5 μl of isopropanol and glycoblue (Invitrogen) diluted 1:150 as co-precipitant. RNA was then dehydrated at −80°C for at least 24-36 h and precipitated by centrifugation at 4°C and maximum speed. Supernatant was removed, the pellet washed once with EtOH 70%, air dried and resuspended in 8 μl nuclease-free water. Library preparation followed the SMARTERÒ stranded Total RNA-Seq Kit v2 – Pico Input Mammalian (Takara Bio, Japan). Due to the degraded nature of input material we omitted the fragmentation step, and 16 enrichment cycles were used in the final RNA-seq amplification. Libraries were then eluted in a minimal volume of 12 μl and sequenced on Illumina Next-Seq500, with 75bp single end reads.

### Bioinformatic data analysis – Single cell data

Raw sequencing data were processed using the CellRanger pipeline (10x genomics, Pleasanton, CA), or kallisto^57^ (for indexed scRNAseq). Count tables were loaded into R and further processed using the Seurat R package^58^. We removed all cells with less than 500 distinct genes observed, or cells with more than 5% of UMIs stemming from mitochondrial genes. PCA was then performed on significantly variable genes, and the first 16 PCs were selected as input for clustering and t-SNE, based on manual inspection of a PC variance plot (“PC elbow plot”). Clustering was performed using the default method from the Seurat package, with resolution parameter set to 5. While lower resolution parameters caused biologically distinct groups with a low number of cells to be merged into single clusters (e.g. sinusoids and arterioles were merged into a single cluster), this relatively large parameter resulted in groups with a high number of cells to be split into an undesirable number of subgroups. We therefore computed the mean scaled gene expression values for each cluster and performed hierarchical clustering of the means using a correlation distance. Clusters with correlations of greater 0.8 were then merged together to result in the final clusters displayed in figure 1b. Marker genes for each population were identified using the FindMarkersAll function and ROC-based test statistics. For mapping of cells to a reference, the Seurat label transfer routines^53^ were used (Figure S6), except in the context of single-cell index-sorting (Figure 6c, S11), where the cell number was insufficient for this method and scmap^49^ was used instead. All mapping results were confirmed by the analysis of marker gene expression (Figure S11b and data not shown). For figure 2e, RNA velocity was run with a neighbourhood size of 50 and a linear velocity scale. The result was insensitive to the choice of parameters, except that the relative arrow lengths varied (data not shown).

### Bioinformatic data analysis – LCM-seq and bulk RNA-seq data

Reads were aligned to version 38.73 of the mouse genome using STAR^59^, and reads falling on exons of genes were counted using htseq-count^60^. Samples containing less than one million reads on feature, as well as two outlier samples identified by PCA, were removed from the data set. Differential expression between niches was determined using the limma/voom workflow^50^ while accounting for batch (i.e. slide number). Cell type proportions in the different samples were then estimated using a custom R implementation of the CIBERSORT algorithm^13^, run on raw count data. CIBERSORT requires the specification of a population-specific gene expression ‘signature matrix’; for that purpose, average gene expression profiles were computed for each cell type and 1571 population-specific genes (Figure S8a, genes were defined by specificity to a given population of 0.8 or greater, as quantified from areas under the ROC curve). To simplify analyses, we merged the highly similar HSPC subtypes into one population for CIBERSORT. Beyond the selection of genes, the CIBERSORT algorithm does not have any free parameters. Further considerations underlying the analyses using CIBERSORT, as well as a discussion on the impact of selecting different sets of genes, are detailed in the Supplementary Note.

For reanalysis of published RNA-seq data (Figure 2c, S4), count tables were created from raw sequencing data available in GEO (GSE89811 and GSE48764), or count tables were downloaded from GEO (GSE109125). For published microarray data, raw expression matrices were downloaded from GEO (GSE33158, GSE43613, GSE57729). It has previously been shown^61^ that CIBERSORT can be reasonably applied to decompose microarray data using a RNA-seq reference.

### Data visualization

All plots were generated using the ggplot2 (v. 3.1.0) and pheatmap (v. 1.0.10) packages in R 3.4.1. Boxplots are defined as follows: The middle line corresponds to the median; lower and upper hinges correspond to first and third quartiles. The upper whisker extends from the hinge to the largest value no further than 1.5 * IQR from the hinge (where IQR is the inter-quartile range, or distance between the first and third quartiles). The lower whisker extends from the hinge to the smallest value at most 1.5 * IQR of the hinge. Data beyond the end of the whiskers are called “outlying” points and are plotted individually^62^.

### RNA-Magnet

We developed RNA-Magnet with the goal of predicting potential physical and signalling interactions between single cells and cell populations, based on expression patterns of cell surface receptors and their cognate binding partners. Potential physical interactions were scored based on receptors that bind to surface molecules expressed on a second cell (e.g. Selectin P ligand-Selectin P, or homophilic interactions of cadherins), or based on receptors binding to structural extracellular matrix components (e.g. Integrin α1β1-Collagen). Signaling interactions were scored based on receptors binding to secreted ligands (e.g. CXCL12-CXCR4).

### Curation of Receptor-Ligand pair lists

This approach depends on reliable lists of ligand-receptor (LR) pairs. Starting from an existing list of human LR pairs^63^, we manually verified all entries for which the mouse orthologues of both the receptor and the ligand were expressed in bone marrow cells. We realized that several entries in this list were not based on intercellular receptor-ligand interactions, but rather intracellular interactions, such as chaperone-receptor interactions (e.g. Calreticulin is not an extracellular ligand but an endoplasmic reticulum bound chaperone of several surface receptors^64^). Other entries were based on mistakes in text mining (e.g. the antibacterial protein Camp was confounded with cAMP), or happened to be co-mentioned in abstracts without evidence for physical binding (e.g. CXCL12 is co-mentioned with CD4 in many abstracts, but we found no support for CXCL12being a ligand of CD4). For some entries, we found no literature reference at all. All of these entries were removed. Furthermore, interactions involving activated components of the complement system were removed as they are irrelevant to homeostatic bone. Ligands were classified as membrane-bound, soluble, or structural ECM components based on Gene Ontology annotations and Uniprot. In ambiguous cases, annotations were verified manually.

The resulting list thereby contained many relevant ligand-receptor pairs, but it lacked most intercellular interactions involving two transmembrane proteins. We therefore used books^65–67^, reviews^68–72^ and the KEGG reactome database to systematically create a list of intercellular interactions involving two transmembrane receptors; literature evidence for each interaction included in the final list is provided in table S3. We paid specific attention to integrin-mediated interactions in order to include information on the specific heterodimers capable of binding a given ligand^68^. In total, the final receptor-ligand pair list contains 721 high-confidence receptor-ligand pairs with expression in bone marrow resident cells (Figure S10a).

### Scoring of receptor and ligand expression in single cells

Surface receptors are frequently expressed lowly at the mRNA level, as we and others have observed in studies combining FACS index sorting and single-cell mRNA sequencing^23,73^. Low efficiencies of reverse transcription (‘dropout’) therefore cause these genes to frequently be missed at the single cell level. While their expression can be imputed using MAGIC^47^, the absolute mRNA level of receptor genes is a poor proxy of absolute amounts of protein (e.g. CD4 mRNA is very lowly, but the protein very highly expressed). We therefore transformed MAGIC estimates of gene expression to a fuzzy logic variable to encode if the gene is ‘expressed’ or ‘not expressed’:

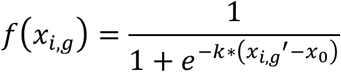

with

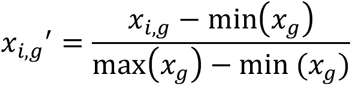

where *x_i,g_* is the MAGIC gene expression estimate of a gene g in cell i. Parameters were set to *x0* = 0.5 and *k* = 10. Here, *x0* specifies the threshold value as a fraction of maximal gene expression above which the gene is considered expressed, and k specifies the degree of fuzziness, with f(x) tending towards Boolean logic for *k* → ∞. Within a reasonable range, the choice of these parameters had no or only a minimal impact on the final result of RNA-Magnet (see Figure S10b).

### Scoring of interaction strength between single cells and reference cell populations

RNA-Magnet provides scores of interaction strengths between a ‘sending’ cell population (i.e. a ligand-secreting cell population) and target ‘receiver’ cells. We therefore first computed population means of gene expression, and then applied a fuzzy logic AND operation to determine if the ligand l is expressed by the sending population K, and the receptor r is expressed by the recipient cell c:

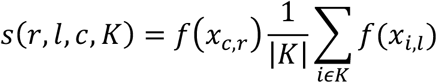

Where |K| is the number of cells in K. The total interaction strength between sending population K and recipient cell c was then computed as

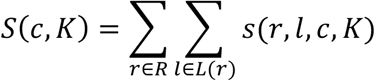

Where R is the set of all receptors and L(r) is the set of all ligands binding to receptor r. In the case of physical (receptor-receptor or receptor-ECM) interactions, we assume that if r is a receptor of l, l is also a receptor of r. *S*(*c, K*) was then normalised to sum to 1 across all populations included.

### Specificity scoring

So far, the interaction score S depends on the total number of ligands secreted by a population K and sensed by a single cell c. This leads to somewhat trivial statements: For example, osteoblasts secrete more ECM components than other populations and the interaction score S between any cell and osteoblasts would therefore be higher than the interaction score between any cell and e.g. sinusoids. In the context of inferring localization, it is therefore crucial to identify if a cell *specifically* interacts with a given cell type. In particular, we observed that highly similar cell types display differential localization (cf. figure 2; for example, fibroblast and CAR subtypes display differential localisation). We account for this by considering the scores relative to an average score seen in similar cells: We computed a local background level of interaction scores for each cell c and sending population K by

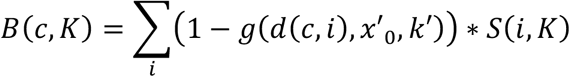

With *d*(*c, i*) specifying the correlation distance between cell c and cell i, and g being a logistic function with parameters x0’ and k’. *B*(*c, K*) was then normalized to sum to 1 across all populations included, and subtracted from *S*(*c, K*) to obtain specificity scores

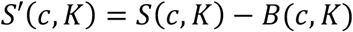

Thereby, we quantified how much more likely a given cell is to interact with a target cell population compared to similar cells. Parameters x0’ and k’ define the size of the neighborhood among which cells were considered similar; they were set such that biologically similar populations (e.g. CAR cell subtypes, fibroblasts subtypes, or vessel types) were included as part of the same neighbourhood (see figure S10c).

In the context of signalling, a similar reasoning applies: CAR cells secrete the largest number of ligands and therefore, the interaction score S between almost any cell and CAR cells is highest, occluding more specific interactions. We therefore again compute specificity scores, but set k to 0 (i.e. we simply take the mean of all cells as background), so as to quantify how much more a given cell is affected by signals from a sending population, compared to the ‘typical’ cell.

### High-level analyses

For figure 5, we used osteoblasts, sinusoidal cells, arteriolar cells and smooth muscle cells as ‘anchor’ population with highly specific localisation to endosteal, sinusoidal and arteriolar niches. We then estimated the preferred localisation of each cell c to one of these four niches N as

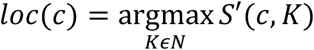

and we estimated an ‘adhesiveness’ score for each cell based on the total number of receptors it expresses (cf. equation 1)

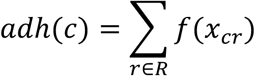

For figure 7a+b, we visualized which populations specifically interact with each other by computing population-wise mean RNA-Magnet scores, and setting a threshold value above which cell types were connected in a graph.

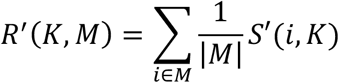

Finally, to obtain an estimate of total signal derived from different niches in figure 7c, we applied RNA-magnet to ligand expression data from LCM-seq.

### Code availability

Our implementation of RNA-Magnet and CIBERSORT, as well as vignettes for re-creating key analysis steps are available at http://git.embl.de/velten/rnamagnet/

### Data availability

Data are available for interactive browsing at http://nicheview.shiny.embl.de. Raw sequencing data and count tables are available through GEO (GSE122467, reviewer access token spqnisgszdopdkh).

**Supplementary Table S4.** Antibodies used in this study.

| Antibody | Clone | Company |
| --- | --- | --- |
| Alpl | Goat Polyclonal | ThermoFisher |
| Anti-Goat IgG AF 546 (whole mount) | Donkey polyclonal | ThermoFisher |
| Anti-Rat IgG DyLight 650 (whole mount) | Donkey polyclonal | ThermoFisher |
| B220 | RA3-6B2 | eBioscience |
| CD105 | MJ7/18 | eBioscience |
| CD106 (VCAM1) | 429 | BioLegend |
| CD11b | M1/70 | eBioscience |
| CD140a (PDGFRa) | APA5 | eBioscience |
| CD144/VE-Cad | VECD1 | BioLegend |
| CD200 | OX-90 | BD |
| CD31 | 390 | eBioscience |
| CD4 | RM4-4 | eBioscience |
| CD41 | eBioMWReg30 | eBioscience |
| CD45 | 30-F11 | eBioscience |
| CD51 | RMV-7 | eBioscience |
| CD61 | 2C9.G2 | BD/BioLegend |
| CD71 | C2 | BD |
| CD8 | 53-6.7 | BD |
| Collagen I | Rabbit polyclonal | Bio Trend |
| CXCL12/SDF-1 | 79018 | R&D |
| Donkey Anti-Goat IgG H&L | Donkey polyclonal | Abcam |
| Donkey Anti-Rabbit IgG H&L | Donkey polyclonal | Abcam |
| Elastin | Rabbit polyclonal | Abcam |
| Endomucin (IF, LCM) | V.7C7 | eBioscience |
| Endomucin (whole mount) | V.7C7 | Stanta Cruz |
| Goat anti-rat IgG | Goat polyclonal | BioLegend |
| Goat Anti-Syrian hamster IgG H&L | Goat polyclonal | Abcam |
| Gr1 | RB6-8C5 | eBioscience |
| Podoplanin | 8.1.1 | BioLegend |
| Sca-1 (IF, LCM, FACS) | D7 | eBioscience |
| Sca-1 (whole mount) | E13-161.7 | BioLegend |
| SM22 | Rabbit polyclonal | Abcam |
| Streptavidin APC-eFluor™ 780<br>Conjugate | - | eBioscience |
| Ter119 | TER-119 | eBioscience |

## Supplementary Note

### Cell type decomposition from spatial transcriptomics using CIBERSORT

CIBERSORT^13^ is an algorithm for estimating the cell type composition of a bulk sample, given a gene expression profile of the sample and a known gene expression profile for each cell type potentially contributing to the sample. Mathematically, the expected expression level x_j_ of gene j in a bulk sample is the sum of cell type averages, s_ij_, weighted by cell type fractions a_i_:

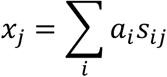

Since the number of genes included is always much larger than the number of cell types, this formulation results in a well-determined system of linear equations. Conventional approaches for its solution however fail to distinguish similar populations and are strongly subjected to experimental noise^74^. CIBERSORT avoids these problems through the use of support vector regression, which has been described to a) internally select an optimal subset of minimally correlated genes, b) penalize each cell type going into the estimate, favoring sparse solutions and c) have a linear penalty function, making it more robust against outliers driven by technical variability.

We used a per-cell type average gene expression matrix defined on 1571 genes with specificity to the individual populations (Figure S8a, genes were defined by specificity to a given population of 0.8 or greater, as quantified from areas under the ROC curve); we will discuss below how the choice of marker gene pre-selection impacts our results. To simplify analyses, we merged the highly similar HSPC subtypes into one population for CIBERSORT. In total, 25 cell types were used for all CIBERSORT analyses.

### Evaluation using simulations and bulk RNA sequencing

To critically evaluate the performance of the CIBERSORT algorithm, we performed a simulation study and confirmed the results using bulk RNA-sequencing. As detailed in the following, we found that the algorithm excels at comparing relative cell type abundancies between niches (i.e. ‘cell type X localizes to niche A over niche B and niche C’), but performs only moderately at estimating cell type proportions within a single niche (i.e. it cannot draw statements like ‘niche A consists to 70% of cell type X and 30% of cell type Y’). We therefore focus our analyses to statements of the first type.

First, we evaluated the ability of CIBERSORT to estimate cell type proportions in a single niche, i.e. a single bulk RNA sequencing sample composed of the cell types described in figure 1b. For this purpose, we *in silico* created a bulk RNA sequencing sample by drawing cell type frequencies from a uniform Dirichlet distribution with 25 dimensions, resulting in a vector of cell type frequencies ***a*** (ground truth). We then assumed that a pooled sample of a total of 1000 cells was to be sequenced. We sampled 1000\****a*** single cells from each population in our main dataset, and summed the gene expression values for each gene across all cells contained in the sample, resulting in a gene expression vector ***x***. This vector was then decomposed using CIBERSORT to result in an estimate of cell type frequencies ***â***. We found that Pearson and Spearman correlations between the ground truth ***a*** and the estimate ***â*** were on the order of 0.6 (Figure S8b, c); however, populations contributing with more than 1% were identified reliably with an area under the curve (AUC) of 0.95. Correlations improved to above 0.9 if a smaller number of cell types were selected that contribute to the bulk sample, while leaving the population reference unchanged (Figure S8c).

To confirm this result, we created bulk RNA sequencing data of total bone marrow and compared the CIBERSORT estimate of its cellular composition to the estimate from our single cell RNA-seq experiment. We found that despite the different RNA-seq protocols used, the performance was as expected from our simulation study (Figure S8d; R=0.71, median correlation for a sample composed of 12 cell types: 0.77).

Next, we evaluated the ability of CIBERSORT to estimate changes in cell type proportions across multiple samples. For this, we repeated the sampling experiment 15 times and quantified the correlation between estimates across samples for each cell type (Figure S8e, f). An optimal performance with correlations >0.95 was found for all populations.

To confirm this result, we used FACS to assemble 8 different pools of B220+ B-cells, CD3+ T/NK-T cells and Gr1+SSC^high^ neutrophils. Each pool contained between 5 and 80 cells of each type, for a total of 100 cells. Pools were then fixed and processed using the same protocol used for the laser microdissected samples, and CIBERSORT was used to quantify their composition. As expected from the simulation study, changes in cell type proportions across samples were very accurately identified with a Pearson R of 0.83-0.92 (Figure S8g).

In line with previous studies^13,75^, these analyses suggest that CIBERSORT excels at identifying changes in cell type proportion across multiple samples but performs only moderately at estimating cell type proportions in a single sample. We therefore restrict the use of CIBERSORT to comparing relative cell type abundancies between niches, and do not determine absolute cell frequencies.

Finally, we also evaluated the extent to which CIBERSORT is capable of discriminating cell types that exhibit similar gene expression profiles (e.g. distinct fibroblast populations or Adipo- and Osteo-CAR cells). We therefore simulated 100 bulk samples assuming that cell types co-occur in a pre-specified manner. Mathematically, we first manually specified a correlation structure of cell type co-occurrence **C** (of dimensions 25×25). We then sampled a cell type frequency matrix **A** (of dimensions 100×25) with a covariance structure **C**. Importantly, correlations were thereby specified and sampled at the level of cell types, and not at the level of genes. We then created a bulk RNA expression profile *in silico* for each sample (row in **A**) as described above, and applied CIBERSORT to estimate its cellular composition. This resulted in a matrix of estimated cell type proportions **Â**. Column correlation structures of **A** and **Â** are compared in figure S8h. Importantly, cell type co-occurrence is correctly identified and not influenced by similarity in the gene expression profile of the reference populations.

In summary, the simulation and bulk RNA sequencing study performed here supports previous evaluations of CIBERSORT: The algorithm is ideally suited for identifying changes in cellular composition between multiple samples. A more detailed analyses of its performance e.g. with regard to noise can be found in ref.^13^.

### Impact of marker gene selection on CIBERSORT results

While CIBERSORT internally selects an optimal set of marker genes, it also requires the pre-specification of a set of reasonably specific markers. To gauge the dependence of CIBERSORT results on marker genes, we repeated the analyses of LCM samples (Figure 2) 60 times, each time using a random subset of 50%, 75% or 90% of the marker genes of each population. For each cell type, we subsequently quantified the fraction of resampling runs that result in the same primary location. The result (Figure S8i) allows us to assess the stability of the CIBERSORT estimates as follows:

For Osteoblasts (n=108 marker genes) and Smooth muscle cells (n=82), any 50% of marker genes can be left out while still allowing unanimous placement of these cells at the endosteum or arteries, respectively.

For Arteriolar fibroblasts (n=54), Arteriolar endothelial cells (n=50) and Fibroblast-Chondrocyte precursors (n=37), any 25% of marker genes can be left out while still allowing unanimous placement of these cells at their respective locations.

For Adipo-CAR cells (n=71), Osteo-CAR cells (n=61), Chondrocytes (n=86) and stromal fibroblasts (n=26), if 25% of marker genes are dropped, this resulted in location swaps in between 10% and 30% of cases. However, the swap was mostly between the primary and a potential secondary location of the cells.

For Sinusoidal endothelial cells (n=33), endosteal fibroblasts (n=69), and MSCs (n=23), estimates depended more strictly on lists of marker genes used. Small numbers of specific markers, elevated intra-population heterogeneity and/or a more ubiquitous localization of these cells may be factors contributing to the estimation uncertainty. For sinusoids and endosteal fibroblasts, we provide further evidence for their localization in figures 2b and 4e, respectively.

## References

1. Ramasamy, S. K. et al. Regulation of Hematopoiesis and Osteogenesis by Blood Vessel-Derived Signals. Annu. Rev. Cell Dev. Biol. 32, 649–675 (2016).

2. Morrison, S. J. & Scadden, D. T. The bone marrow niche for haematopoietic stem cells. Nature 505, 327–334 (2014).

3. Wei, Q. & Frenette, P. S. Niches for Hematopoietic Stem Cells and Their Progeny. Immunity 48, 632–648 (2018).

4. Sugiyama, T., Kohara, H., Noda, M. & Nagasawa, T. Maintenance of the hematopoietic stem cell pool by CXCL12-CXCR4 chemokine signaling in bone marrow stromal cell niches. Immunity 25, 977–88 (2006).

5. Ding, L., Saunders, T. L., Enikolopov, G. & Morrison, S. J. Endothelial and perivascular cells maintain haematopoietic stem cells. Nature 481, 457–62 (2012).

6. Ding, L. & Morrison, S. J. Haematopoietic stem cells and early lymphoid progenitors occupy distinct bone marrow niches. Nature 495, 231–235 (2013).

7. Kunisaki, Y. et al. Arteriolar niches maintain haematopoietic stem cell quiescence. Nature 502, 637–643 (2013).

8. Asada, N. et al. Differential cytokine contributions of perivascular haematopoietic stem cell niches. Nat. Cell Biol. 19, 214–223 (2017).

9. Greenbaum, A. et al. CXCL12 in early mesenchymal progenitors is required for haematopoietic stem-cell maintenance. Nature 495, 227–230 (2013).

10. Méndez-Ferrer, S. et al. Mesenchymal and haematopoietic stem cells form a unique bone marrow niche. Nature 466, 829–34 (2010).

11. Gomariz, A. et al. Quantitative spatial analysis of haematopoiesis-regulating stromal cells in the bone marrow microenvironment by 3D microscopy. Nat. Commun. 9, (2018).

12. Zheng, G. X. Y. et al. Massively parallel digital transcriptional profiling of single cells. Nat. Commun. 8, 14049 (2017).

13. Newman, A. M. et al. Robust enumeration of cell subsets from tissue expression profiles. Nat. Methods 12, 453–457 (2015).

14. Baron, M. et al. A Single-Cell Transcriptomic Map of the Human and Mouse Pancreas Reveals Inter-and Intra-cell Population Structure. Cell Syst. 3, 346–360 (2016).

15. Zhu, L., Lei, J., Klei, L., Devlin, B. & Roeder, K. Semisoft clustering of single-cell data. Proc. Natl. Acad. Sci. 116, 466–471 (2018).

16. Boulais, P. E. et al. The Majority of CD45–Ter119–CD31–Bone Marrow Cell Fraction Is of Hematopoietic Origin and Contains Erythroid and Lymphoid Progenitors. Immunity 0, 627–639 (2018).

17. Coutu, D. L., Kokkaliaris, K. D., Kunz, L. & Schroeder, T. Three-dimensional map of nonhematopoietic bone and bone-marrow cells and molecules. Nat. Biotechnol. 1, (2017).

18. Omatsu, Y. et al. The Essential Functions of Adipo-osteogenic Progenitors as the Hematopoietic Stem and Progenitor Cell Niche. Immunity 33, 387–399 (2010).

19. Zhou, B. O. et al. Bone marrow adipocytes promote the regeneration of stem cells and haematopoiesis by secreting SCF. Nat. Cell Biol. 19, 891–903 (2017).

20. Ono, N. et al. Vasculature-Associated Cells Expressing Nestin in Developing Bones Encompass Early Cells in the Osteoblast and Endothelial Lineage. Dev. Cell 29, 330–339 (2014).

21. Baryawno, N. et al. A Cellular Taxonomy of the Bone Marrow Stroma in Homeostasis and Leukemia. Cell 1–18 (2019). doi:10.1016/j.cell.2019.04.040

22. La Manno, G. et al. RNA velocity of single cells. Nature 560, 494–498 (2018).

23. Velten, L. et al. Human haematopoietic stem cell lineage commitment is a continuous process. Nat. Cell Biol. 19, 271–281 (2017).

24. Karamitros, D. et al. Single-cell analysis reveals the continuum of human lympho-myeloid progenitor cells. Nat. Immunol. (2017). doi:10.1038/s41590-017-0001-2

25. Nestorowa, S. et al. A single-cell resolution map of mouse hematopoietic stem and progenitor cell differentiation. Blood 128, e20–e31 (2016).

26. Tusi, B. K. et al. Population snapshots predict early haematopoietic and erythroid hierarchies. Nature (2018). doi:10.1038/nature25741

27. Tikhonova, A. N. et al. The bone marrow microenvironment at single-cell resolution. Nature (2019). doi:10.1038/s41586-019-1104-8

28. Chen, K. H., Boettiger, A. N., Moffitt, J. R., Wang, S. & Zhuang, X. Spatially resolved, highly multiplexed RNA profiling in single cells. Science (80-.). 348, aaa6090–aaa6090 (2015).

29. Lee, J. H. et al. Highly Multiplexed Subcellular RNA Sequencing in Situ. Science (80-.). 343, 1360–1363 (2014).

30. Stahl, P. L. et al. Visualization and analysis of gene expression in tissue sections by spatial transcriptomics. Science (80-.). 353, 78–82 (2016).

31. Medaglia, C. et al. Spatial reconstruction of immune niches by combining photoactivatable reporters and scRNA-seq. Science (80-.). 358, 1622–1626 (2017).

32. Silberstein, L. et al. Proximity-Based Differential Single-Cell Analysis of the Niche to Identify Stem/Progenitor Cell Regulators. Cell Stem Cell 19, 530–543 (2016).

33. Nichterwitz, S. et al. Laser capture microscopy coupled with Smart-seq2 (LCM-seq) for robust and efficient transcriptomic profiling of mouse and human cells. Nat. Commun. 7, 1–11 (2016).

34. Tamura, S. et al. Podoplanin-positive periarteriolar stromal cells promote megakaryocyte growth and proplatelet formation in mice by CLEC-2. Blood 127, 1701–1710 (2016).

35. Gumbiner, B. M. Cell Adhesion: The Molecular Basis of Tissue Architecture and Morphogenesis. Cell 84, 345–357 (1996).

36. Potente, M. & Mäkinen, T. Vascular heterogeneity and specialization in development and disease. Nat. Rev. Mol. Cell Biol. 18, 477–494 (2017).

37. Xu, C. et al. Stem cell factor is selectively secreted by arterial endothelial cells in bone marrow. Nat. Commun. 9, 2449 (2018).

38. Vento-Tormo, R. et al. Single-cell reconstruction of the early maternal–fetal interface in humans. Nature 563, 347–353 (2018).

39. Camp, J. G. et al. Multilineage communication regulates human liver bud development from pluripotency. Nature 546, 533–538 (2017).

40. Cohen, M. et al. Lung Single-Cell Signaling Interaction Map Reveals Basophil Role in Macrophage Imprinting. Cell 175, 1031–1044.e18 (2018).

41. Siddiqui, J. A. & Partridge, N. C. Physiological Bone Remodeling: Systemic Regulation and Growth Factor Involvement. Physiology 31, 233–245 (2016).

42. Pinho, S. et al. Lineage-Biased Hematopoietic Stem Cells Are Regulated by Distinct Niches. Dev. Cell 44, 634–641.e4 (2018).

43. Itkin, T. et al. Distinct bone marrow blood vessels differentially regulate haematopoiesis. Nature 532, 323–328 (2016).

44. Cordeiro Gomes, A. et al. Hematopoietic Stem Cell Niches Produce Lineage-Instructive Signals to Control Multipotent Progenitor Differentiation. Immunity 45, 1219–1231 (2016).

45. Chan, C. K. F. et al. Identification and specification of the mouse skeletal stem cell. Cell 160, 285–298 (2015).

46. Zhou, B. O., Yue, R., Murphy, M. M., Peyer, J. G. & Morrison, S. J. Leptin-receptor-expressing mesenchymal stromal cells represent the main source of bone formed by adult bone marrow. Cell Stem Cell 15, 154–168 (2014).

47. van Dijk, D. et al. Recovering Gene Interactions from Single-Cell Data Using Data Diffusion. Cell 1–14 (2018). doi:10.1016/j.cell.2018.05.061

48. Moon, K. R. et al. Visualizing Transitions and Structure for High Dimensional Data Exploration. bioRxiv 120378 (2017). doi:10.1101/120378

49. Kiselev, V. Y., Yiu, A. & Hemberg, M. scmap: projection of single-cell RNA-seq data across data sets. Nat. Methods 15, 359–362 (2018).

50. Law, C. W., Chen, Y., Shi, W. & Smyth, G. K. voom: precision weights unlock linear model analysis tools for RNA-seq read counts. Genome Biol. 15, R29 (2014).

51. Shay, T. & Kang, J. Immunological Genome Project and systems immunology. Trends Immunol. 34, 602–609 (2013).

52. Worthley, D. L. et al. Gremlin 1 identifies a skeletal stem cell with bone, cartilage, and reticular stromal potential. Cell 160, 269–284 (2015).

53. Stuart, T. et al. Comprehensive Integration of Single-Cell Data. Cell 177, 1888–1902.e21 (2019).

54. Hennig, B. P. et al. Large-Scale Low-Cost NGS Library Preparation Using a Robust Tn5 Purification and Tagmentation Protocol. G3 (Bethesda). 8, 79–89 (2018).

55. Nichterwitz, S., Benitez, J. A., Hoogstraaten, R., Deng, Q. & Hedlund, E. LCM-Seq: A Method for Spatial Transcriptomic Profiling Using Laser Capture Microdissection Coupled with PolyA-Based RNA Sequencing. in RNA Detection 95–110 (2018). doi:10.1007/978-1-4939-7213-5_6

56. Thomsen, E. R. et al. Fixed single-cell transcriptomic characterization of human radial glial diversity. Nat. Methods 13, 87–93 (2016).

57. Bray, N. L., Pimentel, H., Melsted, P. & Pachter, L. Near-optimal probabilistic RNA-seq quantification. Nat. Biotechnol. 34, 525–527 (2016).

58. Butler, A., Hoffman, P., Smibert, P., Papalexi, E. & Satija, R. Integrating single-cell transcriptomic data across different conditions, technologies, and species. Nat. Biotechnol. 36, 411–420 (2018).

59. Dobin, A. et al. STAR: ultrafast universal RNA-seq aligner. Bioinformatics 29, 15–21 (2013).

60. Anders, S., Pyl, P. T. & Huber, W. HTSeq--a Python framework to work with high-throughput sequencing data. Bioinformatics 31, 166–169 (2015).

61. Chen, B., Khodadoust, M. S., Liu, C. L., Newman, A. M. & Alizadeh, A. A. Profiling Tumor Infiltrating Immune Cells with CIBERSORT. in Cancer Systems Biology (ed. von Stechow, L.) 243–259 (2018). doi:10.1007/978-1-4939-7493-1_12

62. Wickham, H. ggplot2: elegant graphics for data analysis. (Springer New York, 2009).

63. Ramilowski, J. A. et al. A draft network of ligand-receptor-mediated multicellular signalling in human. Nat. Commun. 6, (2015).

64. Orlando, R. A. The low-density lipoprotein receptor-related protein associates with calnexin, calreticulin, and protein disulfide isomerase in receptor-associated-protein-deficient fibroblasts. Exp. Cell Res. 294, 244–53 (2004).

65. Murphy, K. M., Travers, P. & Walport, M. Janeway’s Immunobiology. (Garland Science, 2008).

66. Kreis, T. & Vale, R. D. Guidebook to the extracellular matrix, anchor, and adhesion proteins. (Oxford University Press, 1999).

67. Marks, F., Klingmüller, U. & Müller-Decker, K. Cellular Signal Processing. (Garland Science, 2008).

68. Harburger, D. S. & Calderwood, D. A. Integrin signalling at a glance. J. Cell Sci. 122, 159–163 (2009).

69. Lisabeth, E. M., Falivelli, G. & Pasquale, E. B. Eph receptor signaling and ephrins. Cold Spring Harb. Perspect. Biol. 5, (2013).

70. Sharma, A., Verhaagen, J. & Harvey, A. R. Receptor complexes for each of the Class 3 Semaphorins. Front. Cell. Neurosci. 6, (2012).

71. Miyoshi, J. & Takai, Y. Nectin and nectin-like molecules: biology and pathology. Am. J. Nephrol. 27, 590–604 (2007).

72. Sheppard, D. Integrin-mediated activation of latent transforming growth factor ß. Cancer Metastasis Rev. 24, 395–402 (2005).

73. Paul, F. et al. Transcriptional heterogeneity and lineage commitment in myeloid progenitors. Cell 163, 1663–1677 (2015).

74. Gong, T. et al. Optimal Deconvolution of Transcriptional Profiling Data Using Quadratic Programming with Application to Complex Clinical Blood Samples. PLoS One 6, e27156 (2011).

75. Schelker, M. et al. Estimation of immune cell content in tumour tissue using single-cell RNA-seq data. Nat. Commun. 8, 2032 (2017).

